# Reconstructing the sequence specificities of RNA-binding proteins across eukaryotes

**DOI:** 10.1101/2024.10.15.618476

**Authors:** Alexander Sasse, Debashish Ray, Kaitlin U. Laverty, Cyrus L. Tam, Mihai Albu, Hong Zheng, Olga Lyudovyk, Taykhoom Dalal, Kate Nie, Cedrik Magis, Cedric Notredame, Matthew T. Weirauch, Timothy R. Hughes, Quaid Morris

## Abstract

RNA-binding proteins (RBPs) are key regulators of gene expression. Here, we introduce EuPRI (Eukaryotic Protein-RNA Interactions) – a freely available resource of RNA motifs for 34,736 RBPs from 690 eukaryotes. EuPRI includes *in vitro* binding data for 504 RBPs, including newly collected RNAcompete data for 174 RBPs, along with thousands of reconstructed motifs. We reconstruct these motifs with a new computational platform — Joint Protein-Ligand Embedding (JPLE) — which can detect distant homology relationships and map specificity-determining peptides. EuPRI quadruples the number of known RBP motifs, expanding the motif repertoire across all major eukaryotic clades, and assigning motifs to the majority of human RBPs. EuPRI drastically improves knowledge of RBP motifs in flowering plants. For example, it increases the number of *Arabidopsis thaliana* RBP motifs 7-fold, from 14 to 105. EuPRI also has broad utility for inferring post-transcriptional function and evolutionary relationships. We demonstrate this by predicting a role for 12 *Arabidopsis thaliana* RBPs in RNA stability and identifying rapid and recent evolution of post-transcriptional regulatory networks in worms and plants. In contrast, the vertebrate RNA motif set has remained relatively stable after its drastic expansion between the metazoan and vertebrate ancestors. EuPRI represents a powerful resource for the study of gene regulation across eukaryotes.

## Introduction

RNA-binding proteins (RBPs) bind to transcripts post- and co-transcriptionally to regulate their splicing, poly-adenylation, localization, translation and degradation[1] by recognizing specific RNA sequences, structural elements, or both[2, 3]. These binding specificities can be represented by mathematical models called motifs, which can score RNA sequences based on their likelihood of containing RBP binding sites[4]. Computational motif finding methods[5–12] fit motifs to thousands of bound RNA sequences from large-scale *in vitro* binding assays including RNA Bind-n-Seq[13], RNAcompete[14], and HTR-SELEX[15]. These *in vitro*-derived “intrinsic binding preferences”[4], typically recapitulated *in vivo*[12, 16, 17], are essential for interpreting *in vivo* binding data[18], interpreting non-coding variants[19], and assigning function to RBPs[17]. Some motif models incorporate RNA structure features, but the RNA structure preferences of mRNA-binding proteins are very often well-modelled by a simple lack of base pairing (i.e., nucleotide accessibility) over a short, linear primary sequence motif[5, 12].

Current knowledge of RBP binding preferences is highly biased towards a small number of well-studied RBPs and organisms. Less than 0.1% of all eukaryotic RBPs have any available RNA-binding data, most of which are from mammals or *Drosophila*[15, 17, 18]. RNA motifs can also be assigned to thousands of other RBPs by simple homology rules; RBPs with at least 70% amino acid sequence identity (hereafter abbreviated AA SID) across their RNA-binding domains (RBDs) usually have nearly identical RNA sequence specificities[17]. Most uncharacterized RBPs have less than 70% AA SID to any RBP with an assigned motif, however.

Among eukaryotic RBP families, the RNA recognition motif (RRM) is, by far, the most prevalent sequence-specific RBD and the K-homology domain (KH) is also quite prevalent[2]. The extreme malleability of the sequence specificity of these domains presumably underlies their evolutionary success. Indeed, there are currently almost no cases in which the evolutionary origin of the sequence specificity of extant RBPs in these classes can be traced, or even rationalized.

In principle, “recognition code” homology models – i.e., models that relate the identity of particular residues in an RBD to its RNA sequence specificity[20, 21] – could improve the sensitivity of the “70% rule” described above, as they have for some classes of transcription factors[22, 23]. To be successful, these models require that the modelled RBD class always uses the same set of residues to determine RNA-sequence specificity. Besides the less abundant PUF-domain[20, 21], however, most RBD classes lack these conserved interfaces. For example, RRMs contain two highly conserved RNA-sequence recognition regions, RNP1 and RNP2, but their sequence specificity often depends on residues outside these regions, including linkers and C- and N-terminus domain sequences that recognize specific nucleotides by hydrogen bond interactions[24]. More broadly, the RNA-binding region (RBR) of an RBP commonly contains multiple, adjacent RBDs, and these RBDs can interact with one another to form distinct binding interfaces[25, 26]. This diversity of RNA-recognition interfaces suggests the absence of shared RNA-specificity determining residues for most RBPs[25–27], precluding the use of classic recognition code techniques[22, 23].

A potential solution is training an adaptive “homology model” which learns a similarity metric that predicts shared RNA sequence preferences[23]. One approach, exemplified by the Affinity Regression method[28], is to compute these similarities based on a “peptide profile” which counts short peptide sequences that can be located anywhere within the RBP’s RBR. Affinity Regression assigns weights to each peptide, which are used in computing the similarity between the profiles of two RBRs. Once defined, this similarity measure can be used to infer the RNA sequence preferences of an uncharacterized protein based on the known RNA preferences of similar RBRs. Due to the small number and biased representation of RBPs with known motifs for the adapted similarity metric, only a small fraction of RBPs have been confidently assigned RNA motifs using such methods.

To address these challenges, we generated new binding data for 174 eukaryotic RBPs, and a new algorithm that learns a homology models based on peptide profiles. This new algorithm, Joint Protein-Ligand Embedding (JPLE), uses representation learning[29] within a self-supervised linear auto-encoder framework[30] to adapt its homology model. Combining the new binding data with existing *in vitro* RNAcompete data[17], we used JPLE to identify RNA motifs and RNA-contacting residues for RRM- and KH-domain RBPs across 690 eukaryotes, resulting in motifs for 28,283 RBPs with previously uncharacterized RNA-binding specificities. Combining these motifs with other published datasets and additional inferred motifs, we introduce a resource of 34,737 motifs called EuPRI (Eukaryotic Protein-RNA Interactions), made available via our updated CisBP-RNA web tool: http://cisbp-rna-dev.ccbr.utoronto.ca (*access to be provided on publication*). Using EuPRI and JPLE, we examined the evolution of extant RBP motifs by identifying groups of RRM- and KH-domain containing RBPs with a shared, conserved motif; finding that most RNA motifs appeared recently in multicellular organisms, with clade-specific gain rates, including rapid expansion of motif vocabularies in Nematoda and Angiospermae. Finally, to demonstrate the utility of this resource, we used JPLE assigned motifs to identify 12 RBPs that regulate mRNA stability in *Arabidopsis thaliana*.

## Results

### New RNA motifs for 174 phylogenetically-diverse RBPs

To derive accurate RNA motifs for as many eukaryotic RBPs as possible, we generated new RNA-binding data for RBPs selected to provide useful training data for our homology models while also maximizing the number of RBPs to which we could confidently assign motifs. We selected proteins from 45 well-annotated eukaryotes[31] using a semi-automated procedure to identify potential RBPs containing one or more conventional RBDs that: (i) have many other proteins within 70% AA SID of their putative RBR, i.e., the protein subsequence containing all of the predicted RBDs, (ii) represent both model organisms and under-represented eukaryotic clades, and (iii) would, when combined with pre-existing RNAcompete data[17], provide a more uniform coverage of pairwise protein AA SID levels between measured RBPs. This process produced an initial set of 277 proteins. We measured the intrinsic binding preferences of these candidate RBPs using RNAcompete[17] and identified a subset of 174 RBPs with high quality data using a rigorous, semi-automated quality control procedure[32], thus establishing sequence-specific RNA-binding function for these RBPs. We combined these with pre-existing RNAcompete data[17] to establish a resource of RNA motifs for 379 RBPs (across 381 constructs). This dataset provides broad coverage of RBD architectures and eukaryotic kingdoms, including 41 plant RBPs (**Figures 1A, 1B, S1A, Table S1**).

**Figure 1.**
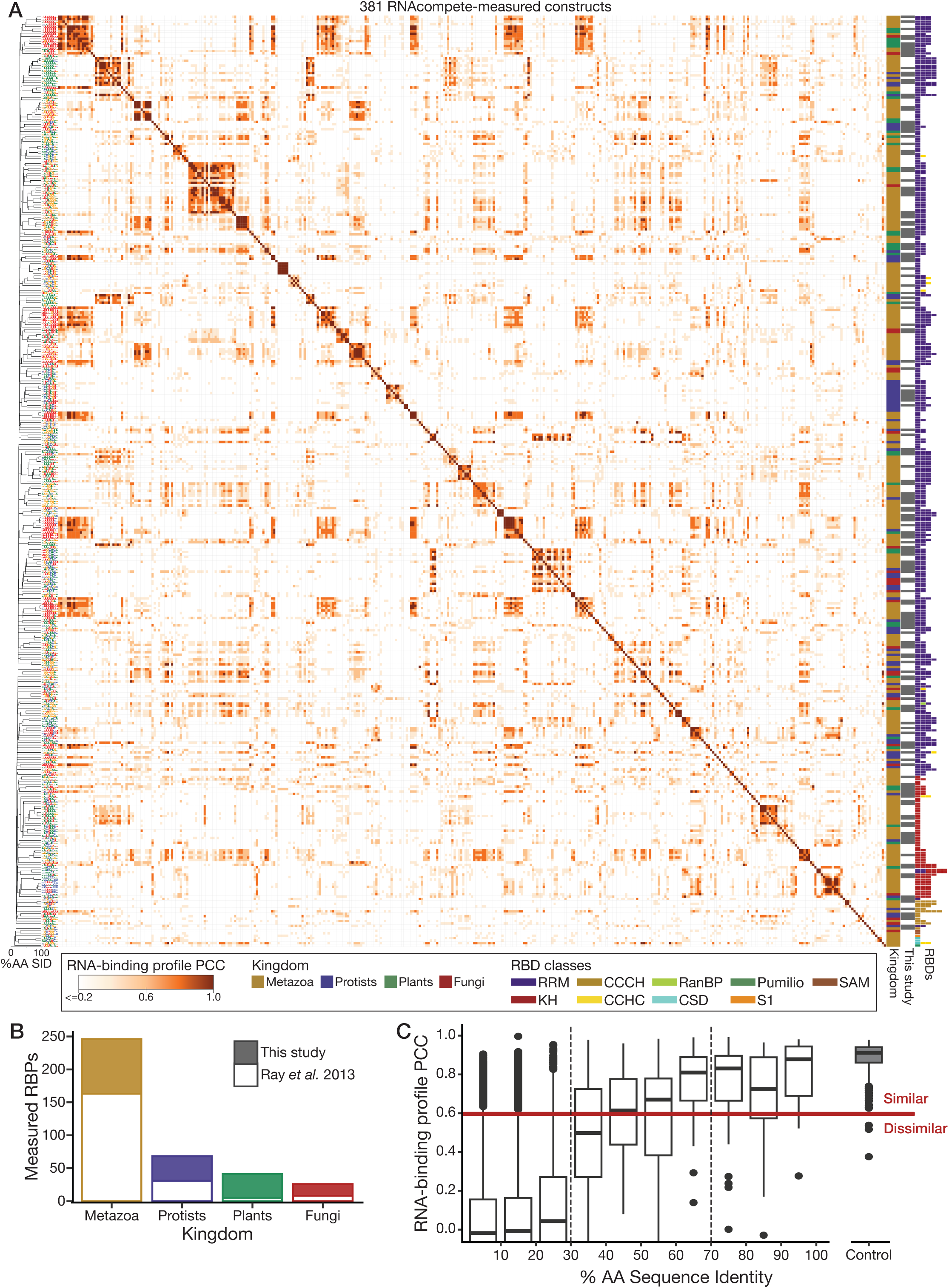
RNAcompete-measured RNA sequence specificities. **A**, Symmetric heatmap displays Pearson correlation coefficients (PCCs) of RNAcompete RNA-binding profiles for each pair of RNAcompete-measured constructs, PCCs < 0.2 are set to 0.2. RBPs are clustered by the AA SID of their RBRs using single linkage hierarchical clustering. Logos of position frequency matrices derived from the top 10 RNAcompete 7-mers are depicted to the left. Indicated to the right is the eukaryotic kingdom for each RBP, whether it is newly measured in this study (grey), and the count and class of RBDs in the construct. **B**, Count of RNAcompete-measured RBPs across eukaryotic kingdoms for RBPs measured for this study and measured previously. **C**, Distribution of RNA-binding profile PCCs for pairs of RNAcompete-measured RBPs whose RBRs fall within the AA SID range indicated on the x-axis. As a control, the distribution of PCCs between RNAcompete Set A and Set B for the same experiment are displayed to the right.

Standard quantification of RNAcompete data produces an estimate of the relative binding affinity (aka “Z-score”) of an RBP to every possible RNA 7-mer[32]. We refer to the vector containing these 4^7^ (= 16,384) Z-scores as the RBP’s *RNA-binding profile* and use it as a detailed representation of the RNA motif implied by the RNAcompete data. We use the Pearson Correlation Coefficient (PCC) of RNA-binding profiles to measure the similarity of two RNA motifs (**Figure 1A**).

Two lines of evidence support the success of our experimental strategy. First, our strategy for choosing diverse RBPs produced a broad range of RNA sequence specificities: the dataset contains 157 distinct clusters of RNA motifs (**Table S1**). In contrast to a previous report[16], the data contain a wide diversity of RNA motifs, particularly among the RRM-containing proteins; nearly half of the clusters (n=74) contain only one RBP (**Figure S1B**) and among the 306 RRM-containing proteins, up to 53% of all possible 7-mers are specifically bound by at least one RBP (**Figure S1C**). Second, the previously reported separation of pairs of RBPs into three classes – highly similar (>70% AA SID, PCC>0.6), variable (30%-70% AA SID, variable PCC) and dissimilar (<30% AA SID, PCC<0.6) pairs – was recapitulated (**Figure 1C**), even when considering RRM and KH domains individually (**Figures S1D, S1E**). Thus, this resource provides a broad coverage of the space of possible RNA targets while demonstrating that the previously reported relationship[17] between protein AA SID and RNA motif similarity holds across eukaryotes.

The RNAcompete data illustrates the challenge of inferring RNA sequence specificity by amino acid sequence homology alone, particularly among the large group of RBP pairs with 30% to 70% AA SID. In this range, AA SID ceases to be a reliable measure of motif similarity; the RBP pairs in this range are nearly equally divided among those with high and low motif similarity (**Figure 1C**).

### The Joint Protein-Ligand Embedding (JPLE) algorithm

Taking advantage of the expanded repertoire of motifs, we sought to improve upon the reconstruction of RNA motifs for uncharacterized RBPs at a greater evolutionary distance, i.e., lower sequence homology. To do this we developed JPLE, a new homology model based on peptide profiles. JPLE captures the association between amino acid sequence and RNA sequence specificity by learning a mapping between: (i) a vector, ***p***, i.e., the *peptide profile* of the RBP, representing the presence, absence, and multiplicity of each short peptide observed in the RBR of the RBP; and (ii) a vector, ***r***, i.e., the *RNA-binding profile*, representing the RNA motif as a table of scores for all possible k-mers. In the current implementation of JPLE, ***p*** consists of entries for all possible 5-mer peptides, with one wildcard character, present within the RBRs of the RNAcompete-measured proteins (**Figure 2A**), while ***r*** consists of the 7-mer Z-scores derived from RNAcompete.

**Figure 2.**
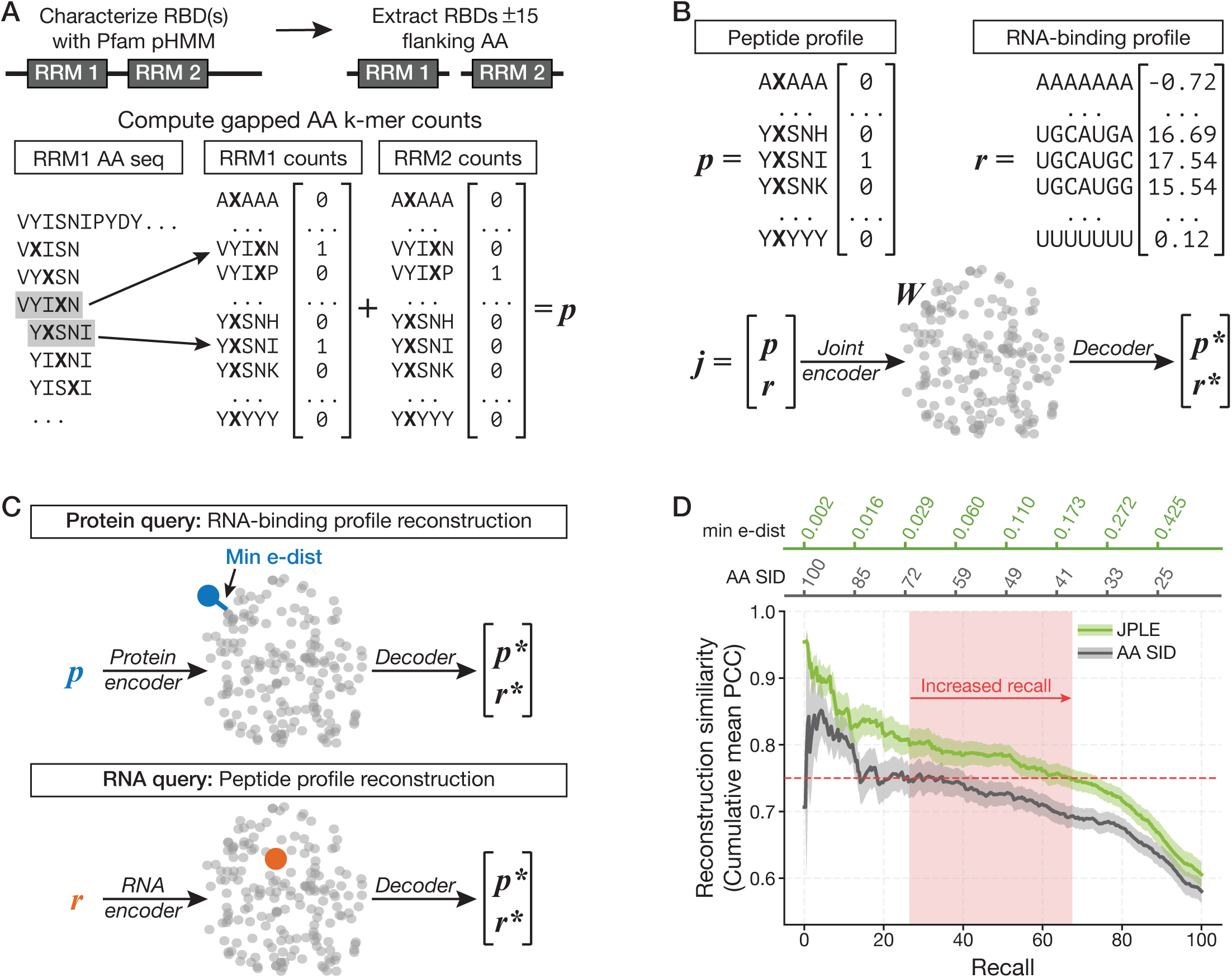
JPLE captures the association between amino acid sequence and RNA sequence specificity. **A**, Derivation of the peptide profile of an RBP. The amino acid sequences of each RBD within the RBP are extracted, along with 15 flanking amino acids, and the occurrence of each amino acid 5-mer with a single wildcard character (X) is counted. An RBP’s peptide profile is the vector of gapped peptide 5-mer counts summed across all its RBDs. **B**, The peptide profile (***p***) is concatenated with the RNA-binding profile (***r***) of an RBP to produce a joint vector (***j***). The joint encoder maps from ***j*** to a low-dimensional embedding, ***W***, and a decoder function maps from ***W*** to reconstructions ***p\**** and ***r\**** of the peptide profile and RNA-binding profile, respectively. **C**, The encoder function can be used with partial input: protein queries (top) estimate ***W*** from ***p*** using a protein-only encoder, and RNA queries (bottom) estimate ***W*** from ***r***, using an RNA-only encoder. Cosine distance in embedding space to the closest RNAcompete-measured RBP embedding (minimum e-dist) is used to assign confidence to this reconstruction. **D**, Precision-recall curves for RNA-binding profile reconstructions generated by AA SID and JPLE. Standard error is shown in the shaded area around each line. Precision (y-axis) is the mean PCC for reconstructions at least as confident as the threshold (top axes). AA SID confidence is AA SID, JPLE confidence is the minimum e-dist. The left boundary of the highlighted region indicates the recall at an AA SID threshold of 70%, at which a mean PCC of 0.75 is achieved. The right boundary of the highlighted region indicates the recall of JPLE at a PCC of 0.75.

In JPLE, the mapping between the peptide profile of a protein, ***p***, and its RNA-binding profile, ***r***, is made using a low-dimensional embedding, ***W***. The embeddings are computed from *joint vectors*, which contain both ***p*** and ***r***, via a *joint encoder* that is trained using a modification of principal component analysis (PCA) (**Figures 2B, S2A**). In standard PCA, a dataset of high-dimensional vectors is transformed into a new, orthogonal, low-dimensional coordinate system (i.e., an embedding) that captures most of the variation in the initial dataset. In JPLE the high-dimensional vectors are the joint vectors derived from the RNAcompete-measured RBPs. In PCA, the new coordinate system is represented by an orthonormal set of high-dimensional vectors, called principal axes, and the coordinates of the low-dimensional embedding of a vector is computed by calculating its projection on each of the principal axes. JPLE’s joint encoder is identical to PCA, except that JPLE’s principal axes are selected only based on their ability to capture the variation in the RNA-binding profile, rather than the joint vector.

Trained on the 355 joint vectors representing each of the 355 RNAcompete-measured RBP constructs that contain only RRM and/or KH domains, JPLE required only 122 axes to explain 96% of the variance in ***R*** (the matrix of all 355 ***r*** vectors), i.e., its low-dimensional embedding space has 122 dimensions (**Figure S2B**). This is a substantial reduction in dimensionality compared to that of each joint vector (n=131,889) and even the number of joint vectors (n=355). By considering only RNA-binding profiles when selecting principal axes, JPLE’s encoder is effectively suppressing the impact, on the embedding, of peptides which have no association with RNA-binding preferences. Overall, JPLE’s embedding only captures 44% of the variance in the peptide profiles (**Figure S2D**), and, as we illustrate below, the captured peptide variations tend to correspond to RNA-contacting peptides that determine RNA specificity.

Once the joint encoder is defined, one can analytically define encoders that optimally estimate the embedding from a subset of the elements in the joint vector (see **Methods**). JPLE defines both a *protein encoder*, that estimates embeddings using only the protein-encoding features in the joint vector, i.e., ***p***, and an *RNA encoder* that uses only the RNA-motif-encoding features, i.e., ***r*** (**Figures 2C, S2C**). Importantly, an embedding computed from any of the encoders can be decoded, reconstructing the high-dimensional joint vector from a weighted combination of the JPLE principal axes, where the weights are derived from the embedding (**Figure 2B**).

Importantly, we found that the similarity between the embeddings of two RBPs is an accurate predictor of the correlation of their RNA-binding profiles (**Figure S2E**). We define the *embedding distance* (*e-dist*) between two RBPs as one minus the cosine similarity of their embeddings, and we use low e-dist as a replacement for high AA SID to determine whether two RBPs have the same RNA motif. The e-dist can be used to assign RNA motifs to uncharacterized RBPs via *protein query* (**Figures 2C, S2C**). This query uses the protein encoder to embed an uncharacterized RBP’s peptide profile, then uses e-dist to identify its nearest neighbors among the embeddings for the RNAcompete-measured RBPs, and finally estimates the RNA-binding profile of the queried RBP as a weighted average of RNA-binding profiles of these nearest neighbors. JPLE therefore provides many of the advantages of traditional PCA: it provides a minimal representation of a dataset, the embeddings themselves are meaningful, and it suppresses non-informative variation (i.e., peptides which do not confer specificity). In subsequent sections, we use JPLE, and the motifs it reconstructs, to address a series of problems in the function and evolution of RBPs and RBP motifs.

### JPLE doubles the number of RBPs assigned RNA motifs

We used a leave-one-out cross-validation framework to assess the reconstruction accuracy of JPLE’s protein queries. **Figure 2D** shows average reconstruction accuracy as a function of increasing e-dist to the closest RBP embedding. Also shown is the average accuracy as a function of increasing AA SID for a simple homology model where the reconstructed RNA-binding profile is that of the training set RBP with the highest AA SID (i.e., its AA SID nearest neighbor). Thresholding the simple homology model at 70% AA SID recapitulates our previous approach for inferring motifs[17]. This cutoff has an average RNA-binding profile reconstruction accuracy of PCC=0.745 and a recall of 27.6%. At an e-dist cutoff of 0.2, JPLE has similar average PCC (=0.748) and a 67.6% recall, a 2.4-fold increase. This gain in recall is equivalent to being able to reconstruct motifs for all held-out RBPs with at least 40% AA SID to an RBP in the embedding (**Figure 2D**, **Table S2**).

To illustrate the value of JPLE’s novel embedding technique, we compared the performance of JPLE to alternative methods with differing protein sequence representations or differing methodology for computing the embedding. First, we evaluated replacing JPLE with linear models trained on fixed-length protein representations derived using the application of modern natural language processing methods to large protein sequence databases[33]. These fixed-length profiles led to substantially lower PCCs between real and reconstructed RNA-binding profiles (**Figures S3A, S3B**, **Table S2**). Next, we evaluated the effect of replacing JPLE’s embeddings with the PCA-based embeddings used by Affinity Regression[28]; this approach had substantially lower recall at the average PCC=0.75 cutoff (**Figure S3C**, **Table S2**). We also evaluated RoseTTAFold2NA[34] and found its ability to differentiate between high and low affinity 7-mers was barely distinguishable from random (**Figure S3D, S3E**). These comparisons underline the importance of JPLE’s strategy to represent protein sequences using peptide profiles, and to learn protein sequence embeddings using joint vectors that include RNA specificity information.

### JPLE predicts RNA specificity-determining residues

The success of JPLE protein queries in predicting RNA motifs suggests that the embeddings for uncharacterized RBPs are encoding the protein sequence features important for target recognition. To investigate what these features are, and how they support RNA motif predictions, we used JPLE RNA queries to reconstruct peptide profiles for known RNA-binding profiles and identified the residues that are represented in the embedding.

Using the RNA encoder, we transformed RNA-binding profiles into the embedding space and reconstructed the peptide profiles (**Figures 2C, S2C**). Because the joint encoder suppresses information about peptides not associated with the RNA-binding profile, and the peptide profiles used to train the joint encoder were counts, we expect the reconstructed peptide profile to contain high values for peptides informative about the RNA specificity of the associated RBP. In practice, because peptides in the profile contain wildcard characters, identifying informative peptides is more complicated, but we can still obtain a *residue importance score*[28] (RIS) for each residue by summing scores from each peptide it appears in. The RIS thus measures the degree to which the identity of a given residue is encoded in the embedding associated with its RNA specificity.

We derived RIS scores for 26 RNAcompete-measured RRM-containing RBPs that had an RRM-RNA co-complex structure in the Protein Data Bank (PDB)[35] (see **Methods**), mapped the RIS values onto the structures, and visualized the result (e.g., **Figures S4A-C**). In general, RIS values were highest for RNA-contacting residues and, often, they were highest for residues that interact with RNA bases rather than the RNA backbone. For example, in human ELAVL1 (aka HuR), high RIS values were observed for RRM1 and RRM2 beta-sheets and the linker region between the RRMs, consistent with structural analyses of ELAVL1/HuR[36] (**Figures 3A, 3B**). For *C. elegans* asd-1, an ortholog of the human RBFOX proteins, JPLE assigns higher scores than conservation to residues within two loops that establish contacts to the first four nucleotides of its preferred UGCAUG motif[37] (**Figure S4A**). Human proteins SNRPA and SRSF1 provide two examples for which RIS values were highest only for those RNA-contacting residues that determine primary RNA sequence specificity. SNRPA, a stem-loop binding protein, was assigned high RIS values for residues contacting single-stranded RNA in the loop, and low RIS values for conserved residues that contact the double-stranded RNA stem (**Figure S4B**). For SRSF1, JPLE assigns high RIS values to the specificity-determining contacts in one of the pseudo-RRM’s alpha-helices[38], and low RIS values to N-terminal residues that contact the RNA backbone (**Figure S4C**).

**Figure 3.**
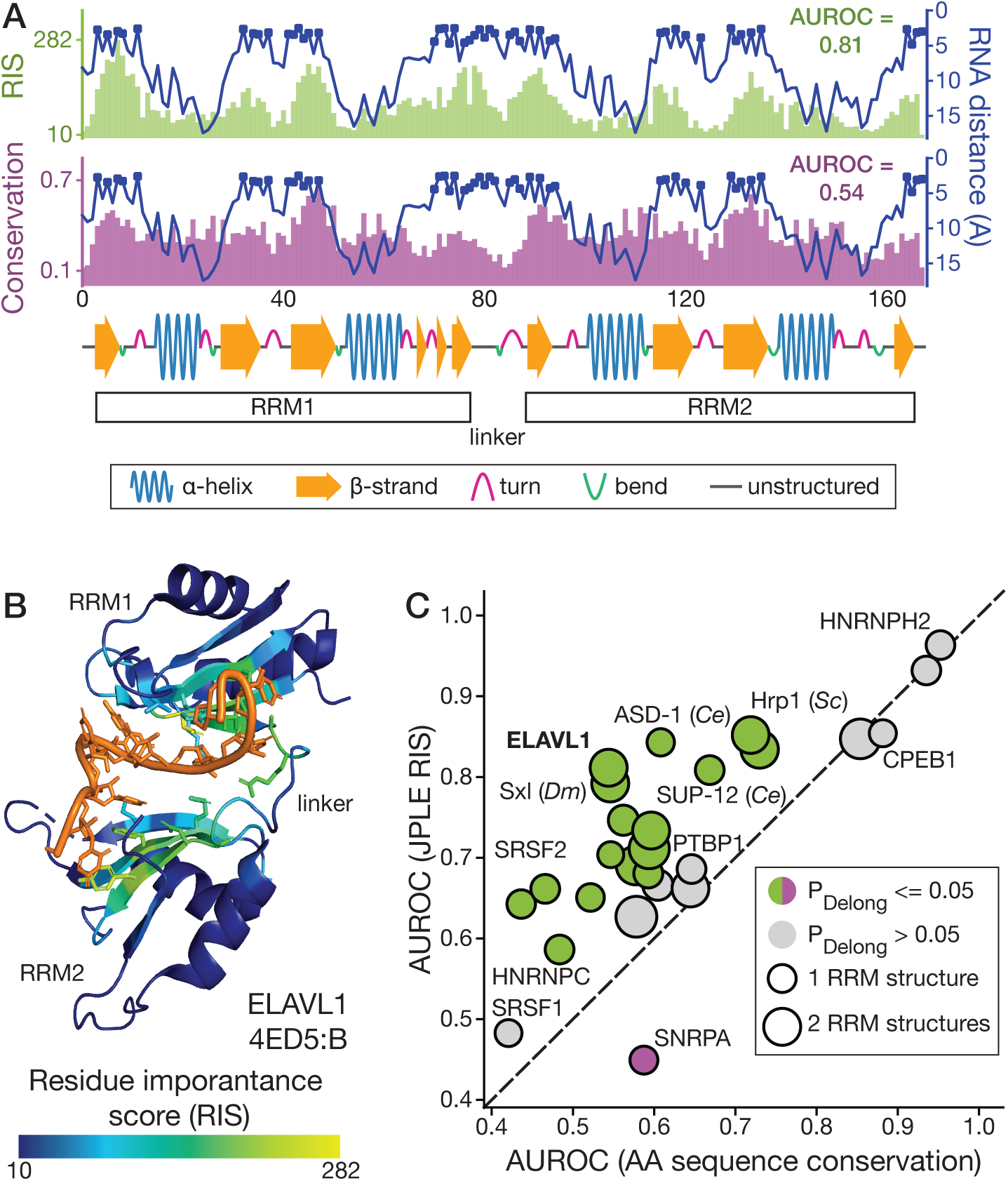
JPLE predicts RNA-interacting amino acids. **A**, The distance between individual amino acid residues and RNA (in Angstroms) is compared to JPLE RIS (top) and conservation scores (middle) for the RBP-RNA co-complex structure in **B**. RNA-contacting residues, (i.e., within 5Å of the RNA) are indicated by dots. A linear visualization of the protein secondary structure is depicted at the bottom. **B**, RBP-RNA co-complex structure (PDB: 4ED5:B) depicts the two N-terminal RRMs from ELAVL1 (*H. sapiens*), with regions colored by JPLE RIS. **C**, Comparison between sequence conservation and JPLE RIS for predicting RRM domain interface residues (i.e., the distance from RNA), evaluated with AUROC. Colored circles indicate a significant difference in performance between the two scoring methods, as determined by the Delong test.

To quantitatively assess these initial observations that JPLE’s RNA queries identify RNA interfaces, we evaluated the ability of RIS values to classify RNA-contacting residues (i.e., those within 5Å of the RNA), using the Area Under the ROC curve (AUROC), and compared their performance to two baselines: sequence conservation and a random forest model trained on the 26 RRM-RNA co-complex crystal structures (see **Methods**). JPLE’s RIS values were better than conservation at identifying RNA-contacting residues for almost all 26 RBPs, and significantly better for 16 (P<0.05, DeLong Test; **Figures 3C, S4D, Table S3**). RIS values were significantly worse only for SNRPA where many of the RNA-contacting residues recognize RNA structure, which is not encoded in the RNA-binding profiles. RIS values had, on average, higher AUROC than the random forest classifier, with significantly better AUROCs for seven RBPs and significantly worse AUROCs for three, including SNRPA (**Figures S4E, S4F**). Collectively, these observations suggest that JPLE’s embeddings are encoding the identity of RNA specificity-determining residues for diverse, and RBP-specific, sets of RBP-RNA interfaces and that JPLE can provide useful structural information for RBPs without known RNA interfaces.

### The EuPRI resource contains motifs for 34,746 RBPs

Next, we took advantage of the fact that JPLE can provide highly accurate reconstructions of RNA motifs for RBPs that are not in the training set. We applied JPLE to assign RNA motifs to as many RBPs as possible among 690 sequenced eukaryotes. For comparison, we also assigned motifs using the 70% rule. We performed protein queries for all KH- and RRM-containing (candidate) RBPs in the predicted proteomes of all 690 genomes **(Figure S5A, Table S4**), and assigned RNA motifs to those whose e-dists to an RNAcompete-measured RBP were below a stringent e-dist cutoff of 0.127. This cutoff guarantees not only an overall average PCC of 0.75 on held-out RBPs, but also a rolling average of PCC>0.70 at all levels of recall up until the cutoff (**Figure S5B**). Of the 76,903 RRM-only RBPs, 10,917 KH-only RBPs, and 307 RBPs that contained both domains, JPLE assigned RNA motifs to 24,320 (32%), 3,749 (34%), and 248 (81%) unmeasured RBPs, respectively (**Figure 4A**), with particularly large gains for plants, fungi, and protists (**Figures 4B, S5C**). As anticipated by our cross-validation studies, the coverage achieved by JPLE is equivalent to being able to assign RNA sequence specificity to every RRM- and KH-domain containing RBP with at least 40% AA SID to an RNAcompete-measured RBP (**Figure S5D**).

**Figure 4.**
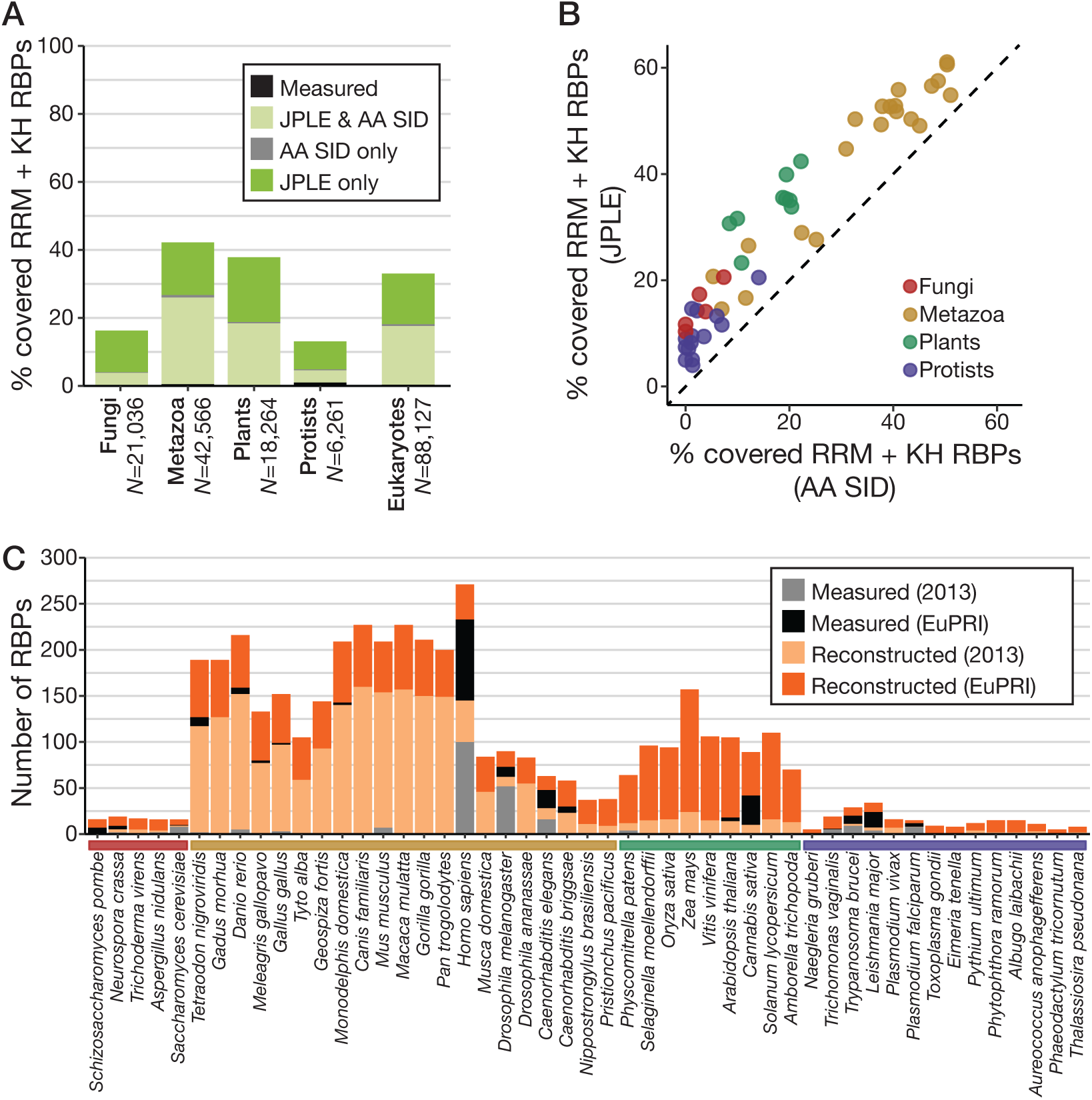
JPLE reconstructs RNA-binding specificities for thousands of eukaryotic RBPs. **A**, The percentage of measured and reconstructed specificities for RRM- and KH-domain RBPs across 690 species is shown across four kingdoms and all eukaryotes combined. The proportion of reconstructed specificities that were identified by AA SID, JPLE, or both are indicated. **B**, Scatterplot displays the percentage of specificities for RRM- and KH-domain RBPs that were reconstructed by JPLE compared to AA SID with the 70%-rule for 49 representative species (listed in panel **C**). **C**, The number of measured and reconstructed RBP specificities for 49 eukaryotes contained in EuPRI. Specificities measured and inferred in the 2013 release of the CisBP-RNA database[17] are differentiated from new motifs.

To generate a comprehensive resource of eukaryotic RNA-binding specificities, we combined the RNAcompete-measured and JPLE reconstructed motifs with motifs reported in other large *in vitro* selection studies[15, 16]. Using the 70% rule against all 504 RBPs with measured motifs, we inferred RNA motifs for a further 5,959 RBPs beyond those described above, the majority of which contain RBDs that lack sufficient training data for JPLE (e.g., the CCCH zinc finger domain). We deposited this resource, called EuPRI (Eukaryotic Protein-RNA Interactions), in our CisBP-RNA database (http://cisbp-rna.ccbr.utoronto.ca).

Between directly measured, reconstructed, and inferred motifs, EuPRI provides sequence specificities for 34,746 eukaryotic RBPs. For the 28,667 RNA sequence specificities reconstructed by JPLE, we further performed JPLE RNA queries, thereby assigning a RIS to each residue in the associated RBR and report these values on CisBP-RNA. Altogether, EuPRI provides specificities for about 33% of RBPs from metazoa, 21% from plants, and 10% from fungi, protists, and algae (**Figure S5E**). For humans, EuPRI provides specificities for 196 RBPs with RRM and KH domains, representing 69.5% of all RBPs with these domains. The largest increase in the absolute number of new motifs is in plants; EuPRI adds, on average, measured or predicted motifs for 111 RBPs per plant species, 114 for Angiosperms (**Figure 4C, Table S4**). EuPRI also covers up to 30% of RBPs for important clades of human parasitic protists including *Leishmania major* and *Trypanosoma brucei*. Altogether, this new resource thus provides an invaluable contribution to the study of post-transcriptional regulation across eukaryotes.

### The age, evolutionary origin, and present-day species distribution of 2,568 conserved RNA motifs

EuPRI’s comprehensive coverage of the eukaryotes, together with JPLE’s accuracy at detecting remote homology, provide a unique opportunity to investigate the evolution of RNA specificity. As such, we next sought to use these resources to estimate the age, and representation in extant species, of conserved RBP motifs.

RBP motifs are thought to be highly constrained and, as such, that new motifs are gained mainly via duplication and neofunctionalization, i.e., one of the duplicated proteins retains the ancestral motif, while the other is released from evolutionary constraint. For example, there are at least six QKI homologs in *C. elegans*. Two, *gld-1* and *asd-2*, share a motif with human QKI, while the other four, each measured by RNAcompete, display subtle variations on the QKI motif (**Figure 5A**). These four QKI homologs are clear cases of duplication and neofunctionalization; they have different measured motifs while co-existing with *asd-2* and *gld-1*, which retain the ancestral QKI motif. However, there are also motifs for which there is no clear evidence for duplication and neofunctionalization; human RBM28 and *C. elegans rbm-28* are labelled as one-to-one orthologs in Uniprot but have distinct motifs (**Figure S6A**). Notably, neither has other close homologs in either organism, strongly suggesting lineage-specific specialization. Therefore, traditional sequence- and gene-tree-based measures of identifying orthologs and shared RBP function do not necessarily identify whether they have a conserved motif.

**Figure 5.**
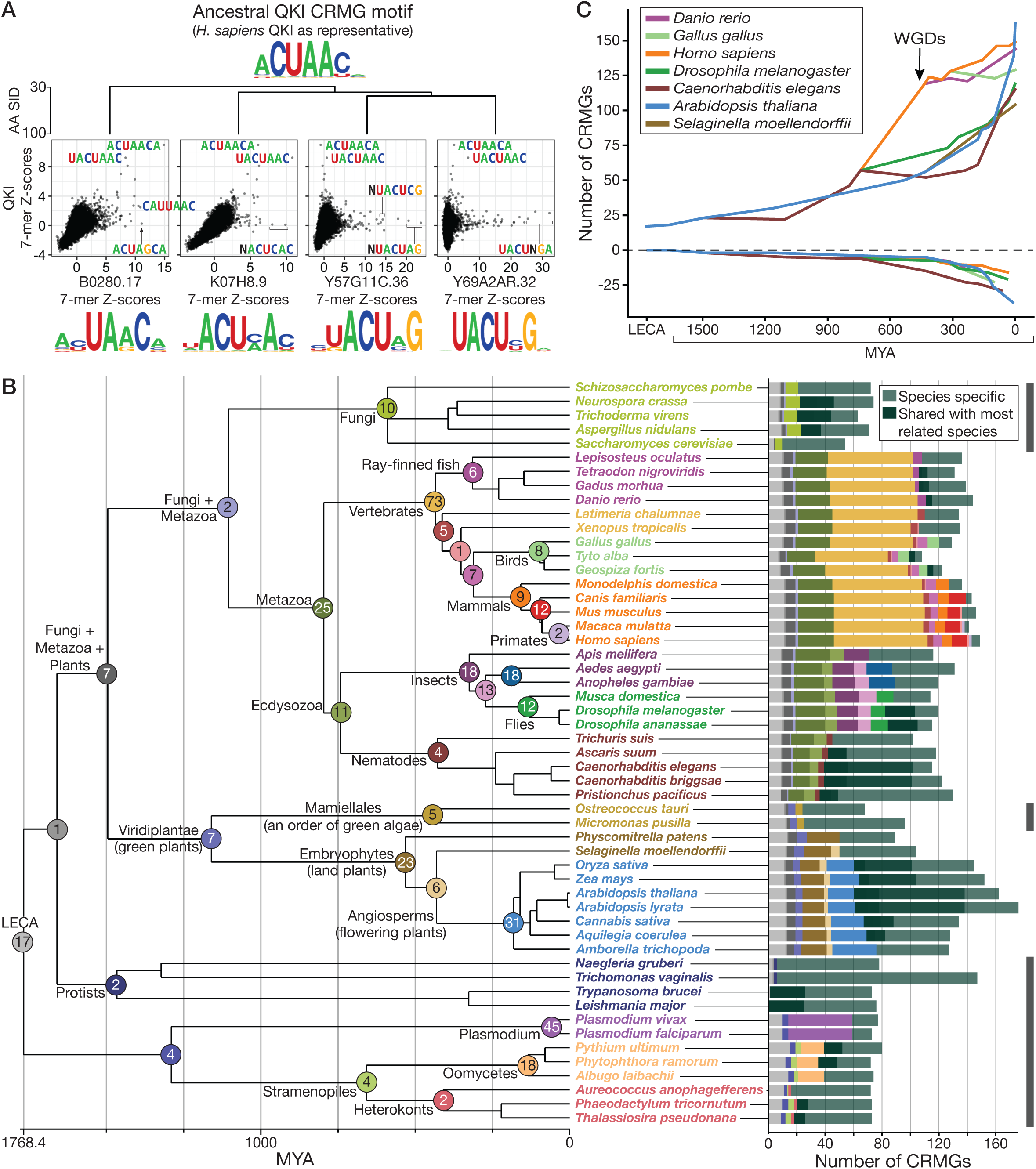
Evolution of eukaryotic CRMGs. **A**, RNAcompete 7-mer Z-scores are compared between QKI (*H. sapiens*) and four homologous *C. elegans* RBPs with diverging binding specificities. Dendrogram above the scatterplots displays the AA SID between the RBRs of the four proteins, the RNAcompete motifs are shown below, and top 7-mers are directly labelled. **B,** (left) Phylogeny of 53 species. Branch points are labeled with the count of new CRMGs gained in the shared ancestor, with branch points of major clades labelled. Font color of species names matches the color of the major recent clade to which they belong. Stacked bar plot counts extant CRMGs in each species broken down by ancestral origin, with colors matching branchpoint colors. Grey bars (far right) indicate unicellular species. **C**, Net number of CRMGs within the common ancestor at different time points is displayed for seven representative species along with the cumulative number of CRMG losses (below zero). The timing of the two whole genome duplications (WGDs) that coincide with the gain of 73 CRMGs in vertebrates is indicated.

To overcome this challenge, we made use of JPLE embeddings, along with traditional methods of identifying orthology, to determine groups of proteins that share a conserved motif. We used parsimony to determine when the motif first appeared, and then used the age of motifs, and their presence in extant species to investigate the large-scale patterns of evolution of eukaryotic RNA motifs.

We selected 53 species that cover the evolutionary space between and within eukaryotic clades and placed all 8,957 RRM- and/or KH-domain RBPs from these species into *conserved RNA motif groups (CRMGs)* (**Table S5**). To generate CRMGs we used e-dist to identify groups of homologous RBPs that have similar motifs, or are within the same e-dist threshold of such groups, using the following criteria: (i) each RBP is grouped with the RBPs to which it has the highest pairwise AA SID; (ii) all RBPs within a CRMG have an observed or predicted RNA-binding profile with PCC>0.6 amongst one another; and (iii) CRMGs are consistent with extant species phylogeny (see **Methods**). The final set of 2,568 CRMGs only includes 831 containing two or more RBPs, 82 of which contain more than 20 RBPs. These outcomes are consistent with the rapid evolution of RNA-binding specificities facilitated by the RRM and KH domains and provide nearly a thousand CRMGs that can be analyzed in terms of species distribution and evolutionary origin.

The multi-protein CRMGs often span distantly related species, indicating that they have ancient origins. To study these origins, we identify the clade and associated ancestral node for each clade by identifying the most recent common ancestor of all extant species with RBPs in the CRMG. **Figure 5B** shows the number of CRMGs in each extant species, the reconstructed ancestral origins of those CRMGs, and, for each ancestor node, the number of CRMGs assigned to it (**Table S6**).

Several observations emerged from this analysis. First, the size of the RNA motif “vocabulary” differs considerably between single- and multi-cellular organisms. Most single-cell organisms in **Figure 5B** had between 60 and 80 CRMGs, whereas almost all multi-cellular organisms had more than 100 and, except for some birds, all vertebrates and flowering plants had more than 125 CRMGs per species. There are some exceptions: *Trichomonas vaginalis*, which has the largest genome among protists[39, 40], had 150 CRMGs among its 183 RRM- and KH-domain RBPs, and *Physcomitrella patens*, a moss, only had 89 CRMGs. Additionally, the genomes of multicellular organisms tend to have more paralogous RBPs in the same CRMG, supporting cell-type-specific functions[41], whereas unicellular organisms generally only have one RBP per CRMG (**Figure S6B, S6C**). Thus, the increased numbers of RBPs in multicellular genomes likely reflect not only cell-type-specific functions of RBPs binding the same motif but also a larger motif vocabulary.

Second, some CRMGs have very ancient origins. Seventeen CRMGs have the last eukaryotic common ancestor (LECA) as their ancestral origin; and 19 CRMGs are present in both plants and metazoa (**Table S6**). These 19 near-universal CRMGs include those representing well-studied orthologous groups of splicing factors represented in humans, e.g., SRSF1, SRSF2, SNRPA, SNRNP70 (with SNRNP35), SF3B6; and CRMGs for each of the nuclear and cytoplasmic poly(A)-binding proteins, PABCP1 and PABPN1. Other near-universal CRMGs contain well-known multi-functional human RBPs, e.g., CELF1 through CELF6 (also known as BRUNOL or CUGBP proteins) are contained in the same CRMG; and RBPs that are members of protein complexes underlying other key post-transcriptional regulatory functions, e.g., mRNA deadenylation by CCR-NOT4 (CNOT4); ribosome assembly (KRR1); the exon-junction complex (RBM8A); and the cleavage and polyadenylation complex (CSTF2).

We also identified several historic periods of rapid motif gain. **Figure 5C** tracks the loss and net gain of CRMGs along specific lineages. The most prominent period of motif growth established 73 new CRMGs and coincided with two whole-genome duplications that occurred between the metazoan and vertebrate ancestors[42]. Among these vertebrate-specific CRMGs are those containing human RBPs HNRNPD, SYNCRIP, and SRSF10. More modest growth in CRMGs occurred between the metazoan ancestor and the shared ancestor with fungi, when 25 new CRMGs were established. Human RBPs in these metazoan-specific CRMGs include PTBP1, QKI, and MSI1. Large motif gains also occurred in the last 200 million years in two clades: nematodes (*Nematoda,* e.g., *C. elegans*) and flowering plants (*Angiospermae*, e.g., *A. thaliana*) (**Figures 5B, 5C**). In both, more than half of the CRMGs in their extant species were established in the last 200 million years, in contrast, for example, to more modest, recent CRMG gains in vertebrates (**Figures 5B, 5C**).

In nematodes, net growth in CRMGs appears to be due to rapid divergence of motifs in all nematodes (i.e., separately in different nematode lineages), coupled with continuous loss of motifs from the metazoan ancestor (**Figure 5C**). Such rapid rewriting and motif gain may reflect an exceptionally high spontaneous rate of gene duplications in, e.g., *C. elegans*[43, 44]. Consistent with our earlier observations, QKI homologs *gld-1* and *asd-2* are members of the QKI CRMG, while the other four homologs are members of four separate CRMGs. In each of the four cases, the ancestral species of their CRMGs is the *Caenorhabditis* ancestor, and each CRMG has the highest pairwise sequence homology with the QKI CRMG, among all the other CRMGs.

In *Angiospermae*, the rapid gains come at the end of a continuous, accelerating gain of motifs. In this lineage, there has been continuous net growth in motif vocabularies from the ancestor of algae and land plants (23 new CRMGs) and from the land plant ancestor to flowering plant (31 new CRMGs). These events occurred with extremely rapid net growth since the *Angiospermae* ancestor, coupled with a relatively high rate of motif loss, suggesting both rapid expansion and rewriting of their post-transcriptional *trans*-regulatory network[45]. For example, *A. thaliana* lost 27 CRMGs, and gained 102 new CRMGs since the *Angiospermae* ancestor (**Figure 5C**). This expansion of motifs mirrors the massive expansion of pentatricopeptide repeat (PPR)-containing RBPs in land plants related to a large increase in complexity of RNA regulation and metabolism[46].

### Using imputed RNA specificities to define the RBP-mediated RNA stability network in *Arabidopsis thaliana*

Taking advantage of these new sequence specificities in plant RBPs, we investigated their functions in post-transcriptional regulation. Previously, we predicted likely regulators of mRNA stability based on whether their motif matches correlate with regulatory patterns present in tissue- and cell-specific RNA expression data[17, 47, 48]. Here, we performed an expanded version of this analysis in *Arabidopsis thaliana*, using its new repertoire of 101 RBPs with assigned motifs and recent expression profiling datasets, as well as, data associating the presence of individual RNA 6-mers in *A. thaliana* transcripts with their half-life in an *A. thaliana* cell line[49].

To estimate a proxy of relative mRNA decay rate for each transcript in each of the 69 RNA-seq-profiled *A. thaliana* tissue types[50], we adapted an algorithm[47] that compares exon and intron read counts. We used these values to generate stability profiles (i.e., the inverse of the mRNA decay rate across all tissue types) for the 6,794 *A. thaliana* genes with reproducible decay rates across replicates (see **Methods; Figure S7A**). For each RBP, we computed the correlation between its expression profile and the stability profile of each mRNA. We then split the set of correlations for a given RBP into two subsets: correlations with predicted targets (mRNAs which contain the RBP motif in the 3’UTR) (see **Methods**), and correlations with predicted non-targets (i.e., the rest of the mRNAs). Finally, to identify putative regulators of mRNA stability, we tested for significant differences in the distributions of these two subsets, determining whether each RBP stabilized its targets (i.e., had comparatively higher correlations), destabilized its targets, or had no apparent effect on stability.

These analyses predicted transcript-destabilizing effects for four RBPs (RBP1, PABN1, SCL30, AT1G78260) and transcript-stabilizing effects for eight RBPs (CID8, CID9, CID10, CID12, AT1G33680, AT2G25970, AT2G46780, AT5G55550) upon binding to the 3’UTR of their target transcripts (**Figures 6A, S7B**). For each of the twelve RBPs, we determined the best matching 6-mers to their motifs; nine of these 12 6-mers were already associated with significantly longer or shorter mRNA half-lives in an *A. thaliana* cell line, and in all but one case, our predicted impact of RBP binding was consistent with the previously assigned impact of the 6-mer in mRNA half-lives (**Table S7**). Intriguingly, the single exception was one of a pair of RBPs with similar predicted motifs (i.e., UG repeats) and identical “best” 6-mers, UGUGUG, but with opposing predicted effects on their targets: we predict AT2G46780 binding increases stability and UGUGUG is associated with longer half-lives in cell line mRNAs[49], whereas we predict AT1G78260 destabilizes mRNAs. So, the impact of UGUGUG on its transcript’s stability depends on which RBP binds it. Both UGUGUG-binding RBPs are expressed in the root and germinating seeds (**Figure 6B**), but AT1G78260 is most highly expressed late in germination, and AT2G46780 in early germination and in the filaments of mature flowers. We also identified another pair of RBPs, RBP1 and AT5G55550, that have a shared RNA sequence specificity but that we predict have opposing impact on RNA stability. Their shared best 6-mer was just below the significance cutoff in the half-life dataset (**Table S7**).

**Figure 6.**
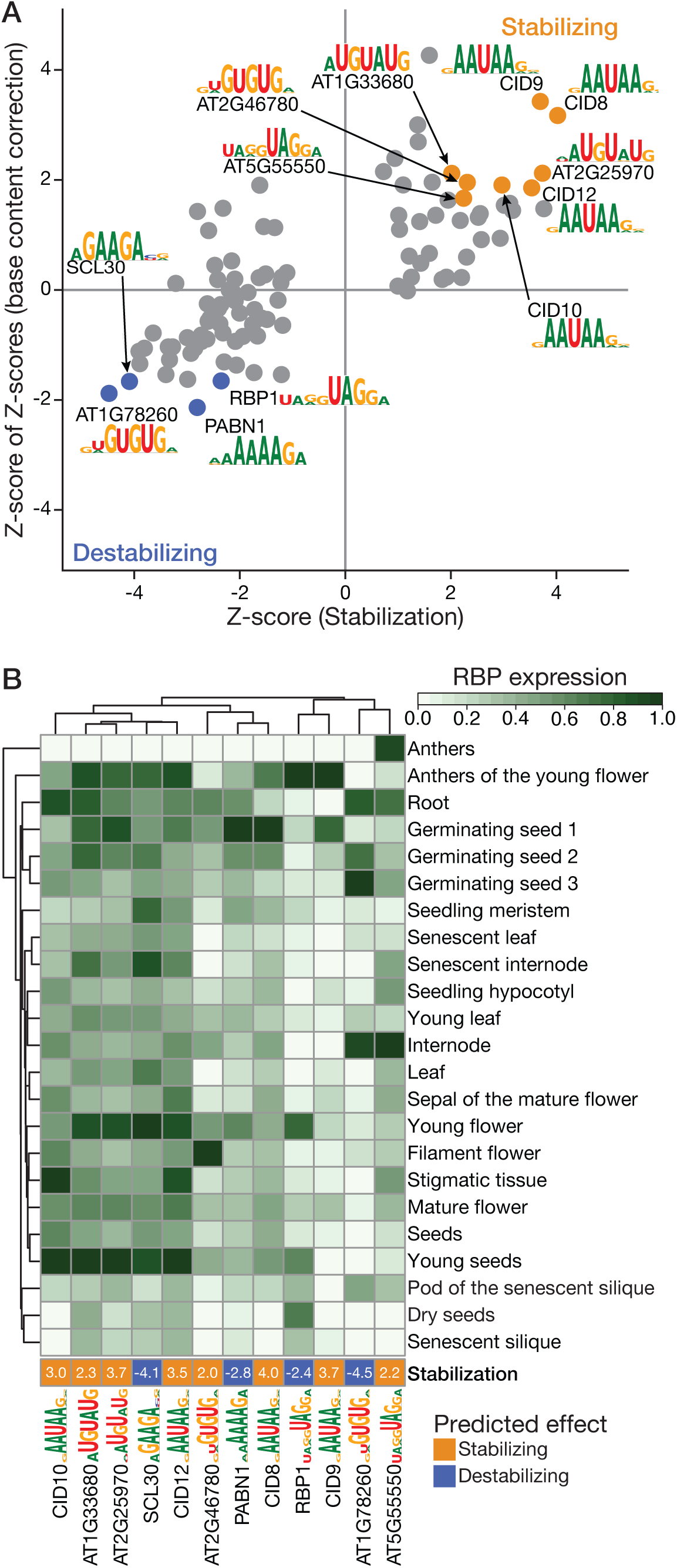
Predicting RNA stability regulators with reconstructed RNA sequence specificities in Arabidopsis thaliana. **A**, RBPs showing significant stabilizing or destabilizing effects on putative mRNA targets (i.e., transcripts containing a 3’UTR motif hit). PCCs are calculated between an RBP’s mRNA expression level and the stability score of each transcript across 69 *A. thaliana* tissues. The distribution of PCCs for putative RBP targets and the distribution of PCCs for non-targets are compared via Mann-Whitney U test to generate a Z-score representative of each RBP’s effect on target stability (x-axis). The y-axis depicts the Z-score of Z-scores which corrects for base content biases by comparing the stabilization Z-score (x-axis) to 100 Z-scores generated from the same test but performed with shuffled RNA sequences. Motifs are displayed for RBPs with Z-scores that indicate a role in regulating mRNA stability (*p*-value < 0.05). **B**, Mean normalized expression level of significant stability regulators across 23 clusters of 69 tissues. Tissue clusters were generated based on tissue mRNA stability profiles (**Figure S7A**). For each RBP, the motif is displayed at the bottom with the stabilization score (Z-score from panel **A**).

In another striking case, all four paralogous CID proteins (CID8, CID9, CID10, CID12) are predicted to stabilize their target transcripts and have partially overlapping expression profiles (**Figure 6B**). These RBPs are named for their CTC-interacting domain (CID), which interacts with the PABC domain (i.e., CTC domain) in poly-A binding proteins (PABPs)[51]. A cleavage and poly-adenylation hexamer (AAUAAA) is one of the most preferred 6-mers for the CID proteins, suggesting they may promote binding of PABPs to transcripts with poly-A tails via CTC domain-mediated protein-protein interactions. Other top 6-mers bound by the CID proteins (GAAUAA, AAUAAG) are enriched in transcripts with long half-lives[49]; these CID RBPs therefore represent potential *trans*-acting factors that contribute to this effect. Overall, our analysis identified trans-acting factors for half-life-associated 6-mers, while also establishing a potential mechanism by which this stabilization occurs and identifying potential tissue-specific regulation.

## Discussion

We have presented a new data resource, EuPRI, and algorithm, JPLE, for studying the RNA-binding preferences of RBPs. EuPRI contains a curated collection of RNAcompete data for 379 RBPs from 33 eukaryotes, including new data for 174 RBPs. JPLE is a computational method that uses low-dimensional embedding to assign RNA motifs based on protein sequence and to identify RNA-specificity-determining residues for RBPs with known motifs. We trained JPLE on the 355 RNAcompete-measured RBP constructs containing only RRM- or KH-domains, but JPLE can be easily expanded to other RBDs if sufficient *in vitro* RNA-binding selection data were available. In principle, JPLE could also be extended to sequence-specific DNA-binding proteins.

Here, we used JPLE to reconstruct RNA-binding specificities for RRM- and KH-domain RBPs across 690 sequenced eukaryotes. Combining this data with other measured and inferred motifs, 23% of eukaryotic RBPs now have an assigned RNA-binding specificity in EuPRI. Plants, and specifically *Angiospermae*, have had the largest increase in RBPs with an assigned motif. Despite being a well-established model organism, only 14 *Arabidopsis thaliana* RBPs had a known motif prior to EuPRI, a number that has now increased by 750%. We were able to assign a function in RNA stability to 12 *A. thaliana* RBPs, nine of their motifs correspond to half-life-associated 6-mers. This is a transformative increase in knowledge about RBP-mediated post-transcriptional regulation in plants.

The success of the JPLE method is noteworthy. JPLE, which can generalize to thousands of RBPs, represents a simple, easily-interpretable linear technique that is a contrast with the currently-popular protein language models based on complex, deep neural network architectures. Protein language models can require significant time investment on expensive and specialized hardware, whereas JPLE runs in seconds on a commodity laptop. Moreover, RNA-specificity-determining residues can be predicted based only on the peptide and RNA 7-mer associations captured by JPLE, suggesting that EuPRI data could be useful for fine-tuning protein language models by providing better features for modelling RBP-RNA complexes. Such a fine-tuning exercise may be critical to assigning RNA specificity to all RBPs; even among the well-studied RRM and KH domain classes, thousands of eukaryotic RBPs could not be confidently assigned an RNA motif by JPLE, whereas in other domains, protein language models may support strong generalization based on limited data.

In addition to its unprecedented scope, EuPRI is qualitatively different from previous datasets. For example, Dominguez and colleagues[16] previously reported a strong bias towards low-complexity motifs among human RBPs, and a tendency for many to bind homopolymeric sequences. In contrast, for the EuPRI dataset, we find much greater diversity in the binding preferences of RBPs, particularly the RRM domain, and a lower prevalence of homopolymers. These differences may be due to the data collection strategy; we focused on assaying widely-conserved, phylogenetically-diverse RBPs, which are more likely to have distinct motifs.

The RNAcompete data illustrates the flexibility of RBP sequence specificity, particularly for the RRM domain. Among these RBPs, RRM proteins collectively show statistically significant binding to nearly half of the possible 7-mers (**Figure S1C**). KH domain RBPs recognize fewer RNA 7-mers, and though there are fewer KH domains across the eukaryotes (and fewer measured by RNAcompete, relative to RRM domains), these trends are consistent with the flexible binding surface of the RRM domain supporting a greater degree of evolutionary innovation in recognized target sequences than the binding cleft used by the KH domains[26, 52]. Perhaps as a result of this flexibility, the RRM domain appears to parallel DNA-binding by the C2H2 zinc finger in terms of divergence of binding sites[23], even between putative one-to-one orthologs, such as human RBM28 and *C. elegans* rbm-28. Surprisingly, 54.3% (143 out of 263) human RRM and KH RBPs have motifs that are younger than the metazoan ancestor, in other words, most human RBPs do not have the same RNA motif as their closest fly or worm homolog; this observation has clear implications in model organism research in PTR. Overall, among the 53 eukaryotes examined, two-thirds of the CRMGs (1,737/2,568) contained only a single protein, indicating that rapid divergence of the sequence specificity of RBPs (most of which contain RRM domains) is very common. This divergence may be due to rapid rewriting of regulatory networks in response to viral or new environmental stresses[53]. Indeed, we see a massive expansion of the number of motifs, and thus the complexity of the post-transcriptional regulatory network, in flowering plants. This parallels the massive expansion of pentatricopeptide repeat (PPR) containing RBPs in land plants; most eukaryotes have <30 PPR proteins, whereas land plant species have >400 (PMID: 24471833). PPR proteins often localize to mitochondria or plastids and it is proposed that they coevolved with organellar RNA metabolism [46, 54]. In contrast, our analysis suggests that at least some of the RRM- and KH-domain containing proteins have cytoplasmic function in RNA stability.

Strikingly, a set of 19 CRMGs shared across plants and metazoa contain RBPs involved in splicing, translation, polyadenylation, and mRNA stability, indicating that these highly conserved regulatory processes are controlled by conserved RNA motifs. One of these near-universal CRMGs contains a rarely studied RBP, RBM42, that interacts with hnRNP K, is a component of stress granules[55], and is MYC-sensitive in lymphoma cells[56]. Three RBPs from this CRMG are measured by RNAcompete: human RBM42, *X. tropicalis* rbm42, and *D. melanogaster* CG2931, all of which bind motifs containing an ACUA core. Orthologous RBPs need not have identical physiological or cellular roles, but it is nonetheless surprising that such a broadly conserved RBP, with conserved RNA-binding activity, would remain so enigmatic.

The EuPRI resource, containing the JPLE predictions, and motif data from other large-scale studies, are consolidated in the CisBP-RNA web server (http://cisbp-rna-dev.ccbr.utoronto.ca (*access to be provided on publication*)). The utility of this resource is exemplified by the prediction of post-transcriptional regulatory functions for *A. thaliana* RBPs, 95% of which had their motifs assigned by JPLE. The same procedure could, in principle, be applied to any sequenced organism with an annotated genome and RNA-seq-based gene expression profiles. This resource will thus be of broad utility in the study of RBPs, inference of post-transcriptional regulatory networks, and prediction of the functional impact of mutations in RBPs and their targets.

## Acknowledgments

This work was supported by a CIHR Foundation Grant to T.R.H. (FDN-148403), a CIHR Project Grant to Q.M. and T.R.H. (PJT-162255), an NIH Grant to T.R.H. (R01 HG008613), and an NIH Grant to Q.M., T.R.H., and M.T.W. (R01 HG013328). Q.M. was partially supported by a NIH/NCI Cancer Center Support Grant (P30 CA008748) and, until 2020, a Canada Artificial Intelligence chair from the Canadian Institute for Advanced Research. T.R.H. is the Billes Chair of Medical Research at the University of Toronto and holds a Canada Research Chair in Decoding Gene Regulation. M.T.W. is supported by NIH Grants (P01 AI150585, P30 AR070549, and U24 HG013078) and a CCHMC CpG Pilot Award (#53553). A.S., K.U.L, and K.N. were partially support by a Vector Institute Research Grant. A.S. was supported by a Mitacs Research Training Award, and K.U.L. was supported by OGS.

## Author Contributions

Conceptualization, Q.M., T.R.H., and M.T.W.; Methodology, Q.M., D.R, A.S., K.U.L., C.N., and C.M.; Software, M.T.W., A.S., M.A., K.U.L, C.L.T., and K.N.; Formal analysis, A.S. K.U.L., C.L.T., O.L., and K.N.; Investigation, A.S., D.R., K.U.L, and H.Z.; Data Curation: M.T.W., A.S. K.U.L., C.L.T., D.R., and M.A.; Writing – Original Draft, Q.M., T.R.H., M.T.W., A.S., and D.R.; Writing – Review & Editing, Q.M., T.R.H., M.T.W., D.R., K.U.L., and C.L.T.; Visualization, A.S. and K.U.L.; Supervision, Q.M., T.R.H., and M.T.W.; Funding Acquisition, Q.M., T.R.H., and M.T.W.; Resources, Q.M., T.R.H., and M.T.W.

## Methods

### Identifying eukaryotic RBPs

To identify RBPs across 690 eukaryotic organisms, we scanned protein sequences for well-characterized RNA-binding domains (RBDs) using HMMER[57] with recommended parameter settings (“Full Sequence E-value” (sequence_eval) <= 0.01 and “Domain conditional E-value” (c-Evalue) <= 0.01). We used RBD Pfam models CSD, KH_1, La, NHL, PUF, S1, SAM_1, YTH, zf-CCCH, zf-CCHC, zf-CCHH, and zf-RanBP from the Pfam database[58]. For RRMs, we defined a new profile HMM (pHMM) as described below.

We grouped all identified RBPs into “RBP families” by their domain architecture, i.e., the type, number, and order of the RBDs in their protein sequence (e.g., RRM, RRM-RRM, KH-RRM, RRM-RRM-KH).

### Generating an extended RRM profile HMM

We extracted all available X-ray and NMR structures of RRMs in complex with RNA from PDB[35] and identified contacts between residues and nucleic acids using COCOMAPS[59]. From these results, we observed that the Pfam RRM_1 pHMM does not include all residues that contact RNA. To generate a new pHMM for the RRM domain, we used the full amino acid sequences of the PDB RRM structures to generate a structure-based (i.e., 3D) multiple sequence alignment (MSA) using the EXPRESSO/3D-COFFEE mode of T-COFFEE with SAP as the structural aligner[60, 61]. To avoid sequence bias, we trimmed the final 3D MSA so that no pair of RRM sequences had an AA SID > 80%. We redefined the limits of the RRM domain to include residues that are in contact with RNA in at least one RRM structure while respecting the secondary structure elements representing the known fold of RRM domains. We used the trimmed 3D MSA to generate a pHMM using HMMER[57], resulting in an extended version of the standard Pfam RRM_1 pHMM that captures 15 additional residues (**File S1**).

### Calculating amino acid sequence identities

We used two separate methodologies for aligning the RNA-binding regions (RBRs) of RBPs and calculating their pairwise amino acid sequence identities (AA SIDs). The first method, “RBP family-wise AA SID calculations”, was used for calculating AA SIDs within RBP families, where all RBPs have the same RBD architecture. We generated an RBR sequence for each protein by concatenating its individual RBD sequences in order and applied clustalOmega[62] with default settings to generate a MSA for all RBR sequences within an RBP family. We defined the AA SID between each pair of RBRs in an RBP family as the proportion of exactly matched, aligned residues in the MSA.

We used a second method for individual pairs of RBPs, “RBP pair-wise AA SID calculations”, to allow for comparisons between RBPs in different RBP families. Here, we defined the RBRs as the subsequence of the RBP that contains all of its RBDs plus up to 15 flanking amino acids before and after each RBD in the RBR. These flanks were added to include linker regions, C- and N-termini. To align two RBRs, we aligned each RBD (with flanks) in one RBR to each RBD in the other RBR using BLOSUM62 scoring (gap opening -11, gap extension -1; Needleman-Wunsch). For each RBD-to-RBD alignment we computed the AA SID (number of exact aligned matches / total length). If the two RBRs had the same number of RBDs (e.g., RRM1-RRM2 and RRM1-RRM2), we calculated the pairwise RBR AA SID as the mean AA SID of each pair of corresponding RRMs (e.g., mean(RRM1 vs. RRM1 AA SID, RRM2 vs. RRM2 AA SID)). If the two RBRs had differing numbers of RBDs (e.g., RRM1-RRM2 and RRM1-RRM2-RRM3), we computed the mean AA SID of all possible alignments of the shorter RBR to the longer RBR, only allowing alignments where adjacent RBDs are aligned (e.g., RRM1-RRM2 can only be aligned to RRM1-RRM2 or RRM2-RRM3 of a three RRM RBR). The maximum AA SID of the shorter RBR alignments is used as the pairwise RBR AA SID. This procedure is designed to account for duplication and deletion events during evolution of protein sequences.

To infer the RNA-binding specificity of an uncharacterized protein based on AA SID (e.g., the “70% rule”), we identified the RNAcompete-measured RBP with the highest overall AA SID and used the RNAcompete RNA-binding profile of this protein as the RNA-binding profile for the uncharacterized protein. The AA SID between the two proteins was used as the confidence score for this prediction.

### Selecting RBPs for RNAcompete

To select eukaryotic RBPs for characterization by RNAcompete, we considered RRM, KH, and CCCH-containing RBPs from 45 well-annotated eukaryotes, representing model organisms and diverse species across the eukaryotic tree[31], identified as described in **Identifying eukaryotic RBPs.** For this task, we used AA SIDs calculated using the RBP family-wise method. We employed four different strategies to select RBPs:

1. To provide sufficient JPLE training data covering various AA similarity ranges, we selected RBPs with differing levels of AA SID to previously characterized RBPs[17]. To this end, we iteratively selected RBPs from the 45 species to evenly populate nine bins of pairwise AA SID (10-19.99% up to 90-99.99%), for each of the RRM and KH RBP families, such that we had at least ten comparisons per RBP family, per bin.
2. We next sought to balance numbers across the major eukaryotic clades. For each major clade (metazoans, plants, algae, fungi, and protists), we selected the single RBP for which the largest number of other RBPs are >70% identical in their RBR. This level was chosen based on the previously established threshold for AA SID-based motif inference[17].
3. To improve sampling of RBPs from diverse model organisms, which we reasoned would be useful to the largest number of investigators, we obtained data for 32 *Cannabis sativa* (plant), 18 *Caenorhabditis elegans* (nematode), 17 *Leishmania major* (parasitic protist), 14 *Homo sapiens* (primate), and 10 *Tetraodon nigroviridis* (fish) RBPs, selected to represent a diversity of RBPs from these species.
4. To generally increase the total number of inferred motifs, we also selected additional uncharacterized RBPs with RBRs that are >70% identical to the largest number of other RBPs, regardless of kingdom.

Collectively, this procedure produced 277 diverse RBPs for experimental characterization using RNAcompete.

### Measuring RNA sequence specificities with RNAcompete

RBP inserts (refer to **Table S1**) were commercially synthesized (Bio Basic) for all 277 selected RBPs and cloned into a custom expression vector, pTH6838, using AscI and SbfI restriction enzymes sites[32]. GST-tagged RBPs were purified from *E. coli* using previously detailed protocols[32] and RNAcompete experiments were performed to determine their RNA-binding profiles.

RNAcompete, a microarray-based *in vitro* RNA-binding assay, has been extensively detailed elsewhere[17, 32]. Briefly, recombinant GST-tagged RBPs are incubated with ∼241,000 designed (not randomized) RNA probes, each about 40 nucleotides long. RNA probes are designed to possess low probabilities for base pairing (i.e., they are single-stranded) and to represent each 9-mer at least 16 times. RBP-RNA complexes are affinity purified and bound RNAs are extracted, labeled with Cy3 or Cy5, and their abundance is measured on a custom Agilent 244K microarray. Measured probe intensities are centered and their variance normalized for each protein (columns) and each RNA probe (rows) to control for variation in RNA and protein concentrations. Finally, RNA sequence specificities are robustly computed as the mean probe intensities of the inner 95% of probes that contained the 7-mer. RNA sequence specificities are transformed to *Z*-scores, setting their mean to zero, and their standard deviation to one. The vector of *Z*-scores for 16,382 7-mers is referred to as the RNA-binding profile; two 7-mers (GCTCTTC and CGAGAAG) are removed because they correspond to the SapI/BspQI restriction site. Position weight matrices are generated by aligning the ten most enriched 7-mers without allowing for gaps. A 7-mer sequence is considered to be “specifically bound” by an RBP if its *Z*-score represents a *p*-value that is equivalent to a Family-wise Error Rate of at most 1% (Bonferroni-corrected *Z*-test).

We evaluated RNAcompete experiments for the 277 selected RBPs according to previously described success criteria[32], and identified a subset of 174 RBPs with successful RNAcompete experiments. Combining these experiments with those from a previous study[17], we generated a set of 420 experiments covering 379 unique RBPs (across 381 unique constructs). For RBPs with more than one RNAcompete experiment, we calculated the mean of the *Z*-scores for the top 10 7-mers and used the experiment with the highest mean in subsequent analyses (**Table S1**).

### Joint Protein-Ligand Embedding

In Joint Protein-Ligand Embedding (JPLE), each RBP is assigned a fixed-length peptide profile vector ***p’***, which contains the count of all 5-mer peptide templates (with four specified amino acids and one wild card) within any of its RBDs including 15 flanking nucleotides on either side of each RBD (**Figure 2A**). This “bag-of-peptides” representation has been used previously for a number of protein function prediction tasks, including homology modelling[28, 63]. Each RBP is also associated with a fixed-length RNA-binding profile vector ***r’*** composed of its RNAcompete *Z*-scores for all RNA 7-mers (**Figure 2B**). The vectors ***p’^T^*** and ***r’^T^*** for the 355 training set RBPs were stacked row-wise to form matrices ***P’*** ∈ ℝ*^n^*^x*p’*^ and ***R’*** ∈ ℝ*^n^*^x*r’*^, where *n* is the number of training set RBPs, *p’* is the number of RBP 5-mers, and *r’* is the number of RNA 7-mers. Columns with zero variance were removed, then each column was centred by subtracting the corresponding column mean (collected in vectors ***μP*** and ***μR*** for ***P*** and ***R***, respectively) and their Euclidean norms were set to one by dividing each column by its norm. The final transformed matrices, ***P*** ∈ ℝ*^n^*^x*p*^ and ***R*** ∈ ℝ*^n^*^x*r*^ have *p* and *r* columns, respectively, and consist of 355 stacked row vectors ***p^T^*** and ***r^T^***, respectively. Each column of ***P*** and ***R*** has zero mean and a norm of one.

The matrices ***P*** and ***R*** were concatenated column-wise to form the joint protein representation ***[P R]*** ∈ ℝ*^n^* ^x^ *^(p+r)^* (**Figure S2A**), to which Singular Value Decomposition (SVD) was applied, giving:

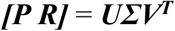

_, where ***U*** ∈ ℝ_*n* x *rank([**P R**])*_, ***Σ*** ∈ ℝ_*rank([**P R**])* x *rank([**P R**])_, **V**_* _∈ ℝ_*(p+r) x rank([**P R**])*_. The diagonal entries of ***Σ***_ are singular values *σi*, whereas the columns of ***U*** and ***V*** (i.e., ***ui***, ***vi***) form the orthonormal basis of ***[P R]*** (***U****^T^**U** = **V**^T^**V*** = ***I****rank([**P R**])*). Typically, when performing dimensionality reduction using SVD, the lowest *σi* are set to zero, effectively removing basis vectors ***ui*** and ***vi*** that explain the least of the variance in ***[P R]***. In JPLE, however, instead of the full matrix ***[P R]***, we retain basis vectors contributing the most to the variance of ***R*** in ***[P R]*** only. To compute the per-basis-vector contribution, we expressed the variance of ***R*** in ***[P R]*** as:

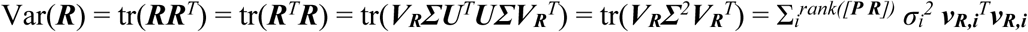

where ***VR*** are the columns of ***V*** which correspond to those of ***R*** in ***[P R]***. The top *d* basis vectors and singular values, ranked by their variance contribution to ***R*** in ***[P R]***, were retained. By performing SVD on ***R*** alone for the 355 training set proteins, we determined that 122 singular vectors were required to achieve a minimum Pearson correlation of 0.95 between their reconstructed (***r*\***) and measured (***r***) RNA-binding profiles (**Figure S2B**). These selected eigenvectors explain 96% of the variance in both ***R*** alone (**Figure S2B**) and ***R*** in ***[P R]*** (**Figure S2D**), demonstrating that *d* = 122 is sufficient to capture most of the variance of ***R*** in ***[P R]***. With this, the original SVD formulation can be written as the following approximation:

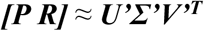

where ***U’*** ∈ ℝ*^n^* ^x^ *^d^*, ***Σ’*** ∈ ℝ*^d^* ^x^ *^d^, **V’*** ∈ ℝ*^(p+r)^ ^x^ ^d^*. The rows of the matrix ***W = U’Σ’*** ∈ ℝ*^n^* ^x^ *^d^* each represent a latent embedding of one of the training set RBPs in the subspace spanned by the columns of ***V’***.

### Protein query

In a protein query, JPLE maps a peptide profile ***p*** to its reconstructed RNA-binding profile ***r*\*** (**Figures 2C, S2C**). Given an RBP with an uncharacterized RNA specificity, JPLE computes its latent embedding ***wu*** by deconvolving its peptide profile ***pu*** according to a mixture matrix containing the orthonormal bases in ***VP’*** (columns of ***V’*** which correspond to those of ***P*** in ***[P R]***), i.e.,

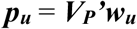

The best approximation ***wu\**** to the embedding of its joint vector is found using Ordinary Least Squares (OLS) regression:

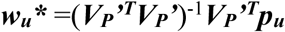

The approximated latent embedding ***wu\**** of the uncharacterized RBP can be mapped to its reconstructed RNA-binding profile ***r*u\*** using one of two different approaches. The first approach, termed global decoding, uses a linear mapping:

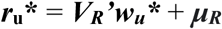

The second approach, termed local decoding, reconstructs the RNA-binding profile based on those training set RBPs with nearby embeddings, thereby implementing a nonlinear mapping. To do so, we first define the e-dist *εui* between an uncharacterized and training set RBP to be the cosine distance between their latent embeddings, i.e.:

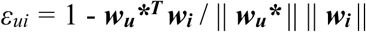

where ***wi*** (i-th row of ***W*** as a column vector) is the latent embedding of the training set RBPi. The e-dist around each uncharacterized RBPu was used with a Radial Basis Function (RBF) kernel (zero mean and *γ* = 25) to obtain the e-sim *εui\**:

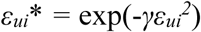

For each uncharacterized RBPu, we constructed a set of neighborhood training set proteins *N* = { *i* | *εui*\* ≥ 0.01 }. The reconstructed RNA-binding profile ***r*u\*** of the uncharacterized RBPu was computed as the average RNA-binding profile ***ri*** of all neighborhood training set proteins, weighted by their e-sim *εui\**:

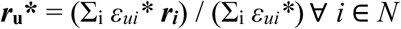

If | *N* | = 0, ***r*u\*** is set to a zero vector. Overall, the local decoder increases the impact of training set proteins with similar embeddings on the reconstruction while negating the impact of training set RBPs with more dissimilar embeddings.

### RNA query

In an RNA query, JPLE maps an RNA-binding profile ***r*** to its reconstructed peptide profile ***p\**** (**Figures 2C, S2C**). While the preserved basis vectors in ***V’*** account for 96% of the variance in the RNA specificities ***R***, they only explain 44% of the variance in the protein peptide profiles ***P*** (**Figure S2D**). Thus, they are correlated with at least one of the RNA specificities, thereby characterizing functionally important peptides for RNA recognition.

To improve the representation of peptide sequences in the training set RBPs, we employed a “data-augmentation” approach to train a version of JPLE for RNA queries. Specifically, we augmented the training set with RBPs that have high homology to those in the training set. These homologs lack RNAcompete measurements but likely have similar binding specificities to those measured by RNAcompete. This data augmentation strategy improved RNA queries but did not have a clear impact on protein queries, so it was not used for the version of JPLE trained for protein queries.

The added homologous RBPs were identified by using HMMER[57] (E-value <= 1e-15) to align RBP sequences containing RRM or KH domains to those with measured RNA specificities. Those uncharacterized RBPs with AA SIDs ranging from 50% to 99% to any measured RBPs were considered hits (i.e., homologs). For a given training set RBP, its 5-mers counts were computed across the alignments of all hits. The resulting peptide counts, normalized by the number of distinct peptides at a given position, were used as a protein representation vector ***p+*** for the given training set RBP. These augmented representations ***P+*** were then concatenated with the measured RNA sequence specificities ***R+*** to train JPLE, resulting in their latent embeddings ***W+*** and orthonormal bases ***V+’***.

Like a protein query, test RBPs were embedded into the latent space ***W+***. However, the latent embedding ***w+u*** of the test RBP was obtained by deconvolving its RNA specificities ru using the mixture matrix V+R’:

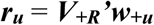

Following this, global decoding was used to map the OLS-approximated latent embedding ***w+u\**** to its reconstructed peptide profile ***p*u\***, representing the relative contribution of each peptide 5-mer to the input RNA specificities:

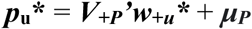

### Alternative JPLE implementations and RNA-binding specificity prediction methods

We retrieved pretrained “Unirep” and “Transformer” (also referred to as “bert” in the code repository) models re-implemented and trained as part of the Tasks Assessing Protein Embeddings (TAPE) framework[33] and embedded the RBR sequences for the 355 training set RBPs using TAPE’s tape-embed command. We then used the 1900- and 768-dimensional embeddings generated by the Unirep and Transformer models, respectively, to train two linear regression models with ridge regression (lambda = 0.0001) to predict the RNA-binding profiles.

Similar to JPLE, Affinity regression (AR) is a machine-learning approach designed for predicting RNA specificities of RBPs[28]. Instead of modeling the direct mapping between ***P*** and ***R***, however, AR learns the interaction, ***A***, between RBP amino acid 4-mer counts ***P*** and RNA 5-mer counts ***D*** to reconstruct ***R*** during training:

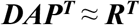

where ***D*** ∈ ℝ*^r^* ^x^ *^r’^*, ***A*** ∈ ℝ*^r’^* ^x^ *^p^*, ***P*** ∈ ℝ*^n^* ^x^ *^p^*, ***R*** ∈ ℝ*^n^* ^x^ *^r^*. We experimented with different formulations of ***D, A,*** and ***P*** and opted for the best performer, where *r* is the number of RNAcompete RNA 7-mers, *r’* is the number of all unique sub-[1–5]-mers within the RNA 7-mers, *p* is the number of RBP amino acid 4-mers, and *n* is the number of training set RBPs. The binary matrix ***D*** indicates whether a given sub-[1–5]-mer is present in each RNA 7-mer. Instead of optimizing ***A*** on the above equation, both sides are multiplied by R, enabling AR to learn the similarity between the RNA specificities of RBPs, i.e.,

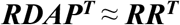

Like JPLE, AR employs SVD to denoise data and reduce computational costs. ***RD*** and ***P*** undergo SVD, retaining singular values and vectors contributing to at least 90% and 95% of their respective variances:

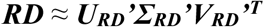

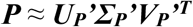

Finally, ***A*** is solved with *L2* regularization:

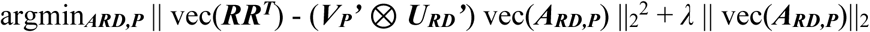

where ***ARD,P*** = ***ΣRD’VRD’^T^A***(***UP’ΣP’***)^T^ and *λ* = 1 / *n*^2^. To reconstruct the RNA-binding profile ***ru****\** of an uncharacterized RBP, the “mapping reconstruction” approach, similar to local decoding in a JPLE protein query, is used by taking the average of the ***r****’s* of all training set RBPs, weighted by the similarity between the training set and uncharacterized RBPs.

For an RBP with uncharacterized RNA specificities, the nearest neighbor model computes the cosine similarity between its peptide profile ***pu*** and all training set peptide profiles ***pi***. The closest training set RBP is then identified:

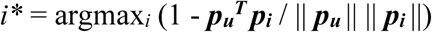

Subsequently, the uncharacterized RBP adopts the RNA-binding profile of the selected training set RBP (***ru\**** = ***ri\****).

Both AR and the nearest neighbor model were applied to the same set of 355 training set RBPs for direct comparison to JPLE.

We evaluated RoseTTAFold2NA’s (RF2NA)[34] capability to differentiate between “binding” and “non-binding” RBP-RNA interactions. For each of the 355 RBPs containing an RRM or KH domain, we generated both a “binding” and “non-binding” set of RNA 7-mers. To do this, we ranked the 7-mers by their Z-scores and excluded the top and bottom 2.5% to account for potential artifacts. The top 50 7-mers were assigned to the “binding” set, while the 50 7-mers closest to the median were designated as the “non-binding” set. This approach produced a total of 35,500 RBP-RNA pairs.

Subsequently, we employed RF2NA to predict the 3D structures of these RBP-RNA pairs. Following the RF2NA evaluation method[34], we used the mean interface predicted aligned error (PAE) as a proxy for RF2NA’s predicted binding specificity, with lower values indicating higher binding specificities. For each RBP-RNA residue pair, we calculated its average PAE by taking the mean of the PAE from the RBP to the RNA residue and the PAE from the RNA to the RBP residue. We then determined the mean interface PAE by averaging these values across all interface residue pairs, defined as RBP-RNA residue pairs within 4.5 angstroms of each other. Structures lacking any interface residue pairs were excluded from the analysis, resulting in 35,479 mean interface PAE values.

### PDB co-complex structures

We retrieved entries of co-complex structures that include both RRM-containing RBPs and RNAs from PDB[35]. These 89 PDB entries contain a total of 156 RBP chains, whose RRM domains were identified and extracted with HMMER[57] using the standard Pfam RRM_1 profile HMM[58]. We identified 156 RRM domains, contained within 119 protein chains, with hmmsearch using default settings[57]. The RRM domain sequences were extended by 35 AAs in both directions and merged if the sequences overlapped in the same protein chain, leading to 118 RRM-containing RBRs.

For each extracted PDB structure, we identified its interface protein residues as those with any carbon atoms within 5Å of any RNA atom. The RBP binding motifs, in IUPAC format, were defined as the connected RNA bases within 5Å of an RBR carbon atom.

To assign RBPs with RNAcompete measurements to those within PDB structures, we computed pairwise AA SIDs and motif overlap. The measured RBP with the highest motif overlap (a minimum of 3.5 nucleotides) and at least 50% AA SID was selected as the best match. Given that most PDB structures have no more than two RRM domains in contact with RNA, some RNAcompete experiments were matched to multiple PDB structures. Subsequently, redundant PDB structures that share more than 70% AA SID were deduplicated by retaining the one with the longest protein sequence.

In total, 27 qualifying PDB structures were assigned a measured RBP; however, we reduced the set to 26 as one PDB structure contained only the third RRM of ELAVL1, which functions primarily as a dimerization domain[64].

### Assignment and assessment of residue importance scores

We conducted RNA queries on all RNAcompete experiments using JPLE with leave-one-out-cross-validation (LOOCV) and obtained their reconstructed peptide profiles ***p*u\**^T^***, which were then stacked row-wise to form the matrix ***P*\*** ∈ ℝ*^n^*^x*p*^. The matrix was standardized by its column mean and standard deviation.

For interface prediction, residue importance scores (RISs) were calculated for each PDB entry from the reconstructed peptide profile ***p*i\*** of the matched RNAcompete-measured RBP. To assign an importance score to each residue in the PDB RBR, we divided each element in ***p*i\*** by its number of occurrences in the RBR, and then, for each residue, we summed the values of the overlapping peptides in ***p*i\***.

As a baseline for interface characterization, we compared JPLE’s RISs to sequence conservation, along with a random forest model trained directly on selected PDB structures. To compute protein sequence conservation, we performed MSA between the RBR sequence of each PDB entry and all RBR sequences from 690 species in CisBP-RNA, and retained those with greater than 30% AA SID to the PDB sequence. We then computed the Jensen-Shannon divergence for each residue in the MSA as a measure for per-residue conservation[65].

For the random forest model, we combined several distinct sequence features: conservation as described above, a position-specific scoring matrix (PSSM) derived from the same protein sequence alignment, physicochemical features, and one-hot amino acid identity[66–68]. The random forest model performed interface predictions for the residue in the centre of a window of five residues. The features of these five residues were summed and provided to the model. The model was trained with LOOCV, focusing only on structures that shared less than 30% AA SID with the tested PDB entry to avoid recalling interface residues from homologous sequences.

To compare the three methods for interface characterization, we computed the Area Under the Receiver Operating Characteristic (AUROC) and the Area Under the Precision-Recall (AUPR) curve over all residues, with residues ranked by their RIS, conservation score, or random forest prediction value.

### Reconstructing RNA sequence specificities for 690 eukaryotes

Using our JPLE model trained on all 355 RNAcompete-measured RBPs, we predicted RNA sequence specificities for RBPs across 690 eukaryotes. RBRs were identified across eukaryotic species as described in **Identifying eukaryotic RBPs**.

We used JPLE to perform protein queries on each of the ∼88,000 detected RBRs with RRM and KH domains. Reconstructions with an e-dist of less than 0.127 were considered confident predictions, corresponding to an average PCC between reconstructed and measured RNA-binding profiles of 0.75 and a rolling average PCC of 0.70 at all levels of recall (**Figure S4B**). Moreover, RNA queries were conducted using the version of JPLE augmented by homologous sequence, as per above, and RISs were computed for all RBRs with confident RNA-binding profile predictions. Secondary structure profiles were computed with SCRATCH[69] for all measured RBR sequences. RBR sequences of unmeasured RBPs were aligned to the measured RBPs, and the predicted secondary structure profile of the RBR with the highest AA SID was used for the visualization on CisBP-RNA.

### Using JPLE e-dist to determine groups of RBRs with common motifs

To study the evolution of RNA motifs in the eukaryotes, we sought to use JPLE to identify groups of RBPs with a conserved RNA motif. First, we investigated the relationship between e-dist and RNA motif similarity for RBPs with low sequence homology to the closest RNAcompete-measured RBP. To do so, we used a neighbor joining algorithm to cluster these proteins into 50 sets of RBPs with high intra-group AA SID and low inter-group AA SID. After clustering, 80% of RBPs had a maximum inter-group AA SID of less than 30%. We trained 50 different JPLEs, each with one of the 50 groups held-out, and evaluated how well e-dist within the held-out group correlated with the PCC between the pairs of held-out, low sequence homology RBPs. We found that pairs of held-out RBPs with e-dist <0.2 had an average PCC of 0.62 (**Figure S2E**), the lower bound of the PCCs of technical replicates (**Figure 1D**). As such, we used an e-dist threshold of 0.2, along with sequence homology to identify clusters of uncharacterized RBPs with a shared motif and derived from the same ancestor. Note that the 0.2 e-dist cutoff is the same as that used in the main text to estimate recall of JPLE in LOOCV, where it is associated with a PCC of 0.748. Here the average correlation is lower because the inference of the RNA profiles of the held-out, low homology RBPs is more challenging.

### Clustering RBPs into Conserved RNA Motif Groups (CRMGs) and characterizing their last common ancestors

To define clusters of evolutionary related RBRs with conserved RNA sequence specificity, we used a multi-step clustering algorithm that incorporated JPLE latent distances, pairwise AA SID, a selected species tree, and agglomerative clustering. We used TimeTree[70] and the ETE Toolkit[71] to select and extract evolutionary distances between 53 eukaryotic species that cover the evolutionary space between and within eukaryotic clades. Collectively these species contain 8,957 RBR sequences that are exclusively composed of RRM and KH domains. We used a global alignment (BLOSUM62 and gap penalty 11,1 (open, extension), Needleman-Wunsch) to compute pairwise AA SIDs between all pairs of RBR sequences (see **Calculating amino acid sequence identities**). Between RBPs with the same or one domain difference, however, we computed sequence identities using the shorter sequence length as a reference to account for homologs that lost a single domain. For all others, we used the longer sequence as a reference, assuming that orthologs generally do not gain or lose more than one domain.

We constructed a “highest homology network” between pairs of RBRs in different species. The “highest homology network” contains one node for each of the 8,957 RBR sequences, and we connected each RBR to the RBR with the highest AA SID in each of the other 52 species. This highest homology network is used as a scaffold for our clustering, thereby ensuring that we only combine RBR sequences likely to be descended from a common ancestral protein into conserved RNA motif groups (CRMGs). We then remove any links in the highest homology network between RBRs with an e-dist of greater than 0.2. This created a set of 2,463 connected components of the filtered network, these components are used as the initial set of CRMGs. Each pair of RBRs in a cluster are thus connected by at least one path where each link is both a highest homology link and has e-dist < 0.2. Examining this initial draft of the CRMGs, we found that some of the initial clusters consisted of two or more groups of highly interconnected RBRs, with the groups connected by a single “transitioning” RBP or one, or a small number of links. We reasoned that these inter-group links were likely false-positives, potentially due to fusions of RBDs from distinct RBPs, and further split the connected component using a procedure that ensured that the resulting CRMGs both were consistent with the phylogeny and had an average e-dist < 0.2 between all pairs of RBRs. To do so, we applied species-tree guided agglomerative clustering (average linkage) on JPLE e-dist, separately to each initial CRMG.

Starting from the two most closely related species in the CRMG, we move the species tree upwards (i.e., further back in time) to the last common ancestor of the CRMG and at each branch point we check if the average e-dist between all descendent RBRs is below 0.2. If at any branch point, the average e-dist is > 0.2, the CRMG is split into two CRMGs. Additionally, we wanted to avoid retention of a CRMG with the highly unlikely evolutionary event that it was lost independently in two or more subsequent branch points. To do so we split a CRMG if a branch point and a subsequent descendent branch each contain a descendant branch that does not contain any RBPs in the CRMG (i.e., it was independently lost twice). This step separated the 2,463 initial clusters into 3,095 *post-split* CRMGs whose e-dists to one another were all > 0.2.

We then merged some post-split clusters in order to combine paralogs within a species that may share RNA-sequence specificity but were not part of the same connected component in the highest homology network (which only connects RBRs between different species). To identify these paralogous groups, we computed the average JPLE distance between all pairs of post-split CRMGs containing RBRs in the same species and merge any post-split CRMGs with average e-dist distance < 0.2. This step reduces the 3,095 post-split CRMGs to 2,866 paralog-merged CRMGs.

Although JPLE is able to correctly distinguish binding similarities between most RBRs, we were able to identify cases of false positives and negative groupings in the paralog-merged CRMGs based on other features. First, single RBRs are removed from a paralog-merged CRMG if removing the RBR changed the common ancestor of the CRMG and if the RBR did not possess a one-to-one triangle orthology connection to any other pair of RBRs in the CRMG. This step generates 14 additional, single RBR groups, thus affecting only 0.5% of clusters. To account for potential false negatives from potentially inaccurate JPLE distances in low-confidence regions (i.e., far from training set RBPs), we combined CRMGs if on average they shared more than 70% AA SID. This step caused the merging of 417 (14.5%) mainly single RBR CRMGs into another CRMG. After this step, we are left with the final version of the CRMGs.

We then assigned each CRMG to a clade represented by the ancestral node in the species tree by inferring the most recent common ancestor of all extant species with RBPs in the CRMG.

### Computing mRNA decay rates from RNA-seq for 69 tissues of Arabidopsis thaliana

We downloaded 138 raw sequencing files from the SRA (Accession: PRJNA314076) corresponding to two RNA-seq replicates for 69 *Arabidopsis thaliana* tissues[50]. We aligned each FASTQ to the TAIR10 genome assembly using Bowtie [72]2 with the -sensitive-local option and filtered for reads with a map quality >= 30 using SAMtools[73]. For each gene, reads that mapped fully within exons were counted separately from reads that fully or partially overlapped with introns. Intronic reads were used as a proxy for pre-mRNA counts. To approximate mature mRNA counts, because exonic reads can originate from both pre-mRNA and mature mRNA, we subtracted the intronic read counts from exonic counts and set small negative values to zero. All count values were normalized to reads per kilobase million.

Steady-state mRNA abundance is determined by transcription, processing, and degradation rate. To eliminate the effect of transcription and processing, we approximated the stability *S* (i.e., the inverse of the mRNA decay rate *kd*) for each gene *j* in tissue *t* from the ratio of exon, *e*, and intron, *i*, counts[47]. In order to compare differential expression of RBPs to the differences of stabilities across tissues downstream, we compute the fold change of the stability to the mean of each gene across tissues. We account for contributions from varying splicing rates through a bias correction that was previously described in[47].

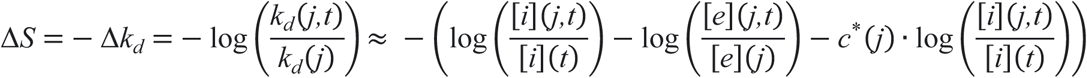

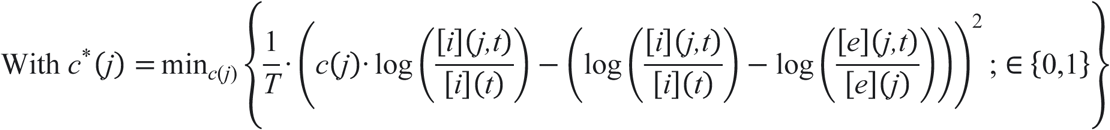

Where *t* is one of the *T* = 69 tissues and [*e*](*j*) and [*i*](*j*) are the mean exon and intron counts of gene *j* across all tissues. We only diverge from this previously-described procedure by restricting the bias constant to be between zero and one, which is implied by its derivation.

Briefly, the gene specific bias constant *c*(*j*) can be derived by approximating the splicing reaction with the Michaelis-Menten equation[47], leading to an extra bias term *b*([*i*](*j*),[*i*](*j,t*),*KM*) that is added to the differential stability *ΔS*:

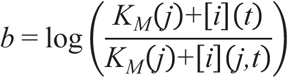

Using Taylor’s approximation for this term around 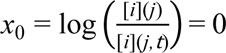 the non-linear bias term can be linearized to:

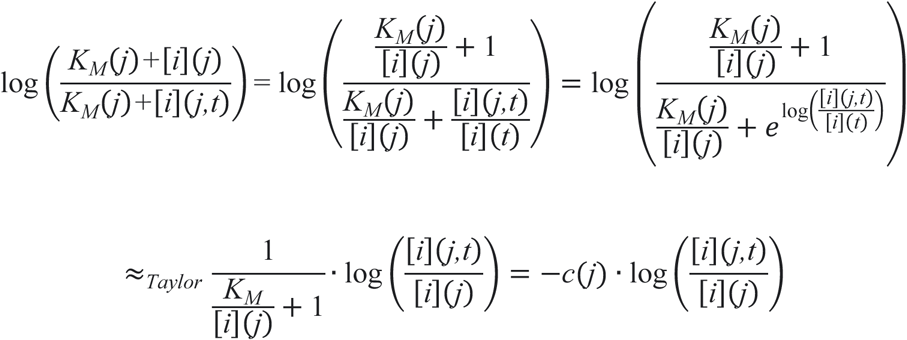

This leads to a bias that linearly scales with 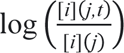 with the bias constant 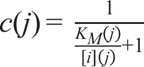 The bias constant *c*(*j*) is dependent on the gene specific Michealis-Menton constant, and the mean intron count for the gene. In accordance with the behaviour of the equation in the extremes, *KM* >> [*i*](*j*) or [*i*](*j*) >> *KM*, *c*(*j*) can only take values between zero and one, where *c*(*j*) ∈ {0,1}. Due to this linear relationship, we approximate the bias *c*(*j*) with gene-specific constrained linear regression with 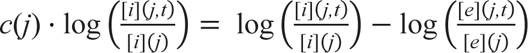. The fitted value *c\**(*j*) is subtracted from the difference between introns and exons for an unbiased estimate of the degradation rate. The solution *c\**(*j*) estimates the value of the bias at which we observe the smallest differential effect on the stability 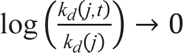, and therefore represent a conservative estimate of the differential stability.

We filtered the set of genes for use in downstream analysis to the 6794 genes that had a Pearson correlation >= 0.6 for intron counts, exon counts, and mRNA stability scores between replicates across all 69 tissues.

### Identification of RBPs with effect on mRNA stability in Arabidopsis thaliana

We extracted *A. thaliana* 3’UTRs sequences and selected the single longest 3’UTR isoform as the representative 3’UTR for each gene. We scanned 3’UTR sequences with motifs for 101 RBPs, derived from five RNAcompete-measured and 96-JPLE reconstructed RBP specificities. Three *A. thaliana* RBPs with inferred specificities were excluded as they could only be reconstructed by AA SID and not by JPLE. First, we transformed the 101 position frequency matrices into position specific affinity matrices (PSAM) by setting their max value at each position to 1. We then slid the PSAMs across the 3′UTR sequences and computed the binding probability at each position as the product of matching base positions along the entire PSAM. Since binding to a sequence is affected by the match of the sequence to the RBPs’ sequence specificity as well as the abundance of putative binding sites within the target, we computed the binding score for each 3′UTR as the sum of all scores along the length of the sequence that had a value greater than 10% of the maximum possible binding score.

RBP sequence specificities enable statistical testing for the effect of *trans*-acting factors on RNA stability, instead of solely testing for the effect of motifs in *cis*. We used computed exonic expression of an RBP as a proxy for the RBP’s protein abundance and computed the correlation between the expression of each RBP and the stability of each transcript across 69 tissues. We then determined RBP-specific binding score thresholds to determine the final high confidence set of putatively bound transcripts. The threshold for 3’UTR to be defined as putatively bound by an RBP was defined as the binding score at which the difference between the correlations between putatively bound and unbound sequences had the lowest *p*-value under a Mann-Whitney U test (two-sided). We then determined the difference of this correlation between transcripts with and without putative binding sites and corrected the computed effect for multiple hypotheses with Benjamini-Hochberg across RBPs (statistic referred to here as *p*-valueFDR). In addition, we randomly shuffled 3’UTRs 100 times to determine if the *Z*-score statistic calculated with the real 3’UTRs is significantly different from *Z*-score statistics calculated with a set of shuffled 3’UTR sequences with the same RBP motif (*t*-test, statistic referred to here as *p*-value*Z*). This statistic corrects for nucleotide or expression biases. RBPs that possess significant effects (i.e., *p*-valueFDR < 0.05 & *p*-value*Z* < 0.05) in both tests were identified as putative stability-regulating RBPs.

### Data Availability

RNAcompete normalized intensity data is available at GEO (http://www.ncbi.nlm.nih.gov/geo/) under accession number (*data to be provided on publication*). Code is available on GitHub: https://github.com/LXsasse/RBPbinding

## Supplemental Tables and Files

**Table S1. RNAcompete experimental details.**

**Table S2. Performance of JPLE and other RNA-specificity prediction methods for the 355 training set proteins.**

**Table S3. Performance of residue importance scores and other prediction metrics for 26 PDB co-complex structures.**

**Table S4. Count of identified RBPs and RBPs with assigned motifs across 690 eukaryotes.**

**Table S5. Conserved RNA motif group assignments for 8,957 RBPs from 53 species.**

**Table S6. Conserved RNA motif group ages and clade assignments.**

**Table S7. Half-life data for putative stability-regulating A. thaliana RBPs.**

**File S1. Extended profile HMM for the RRM domain.**

## Supplemental Figures

**Figure S1.**
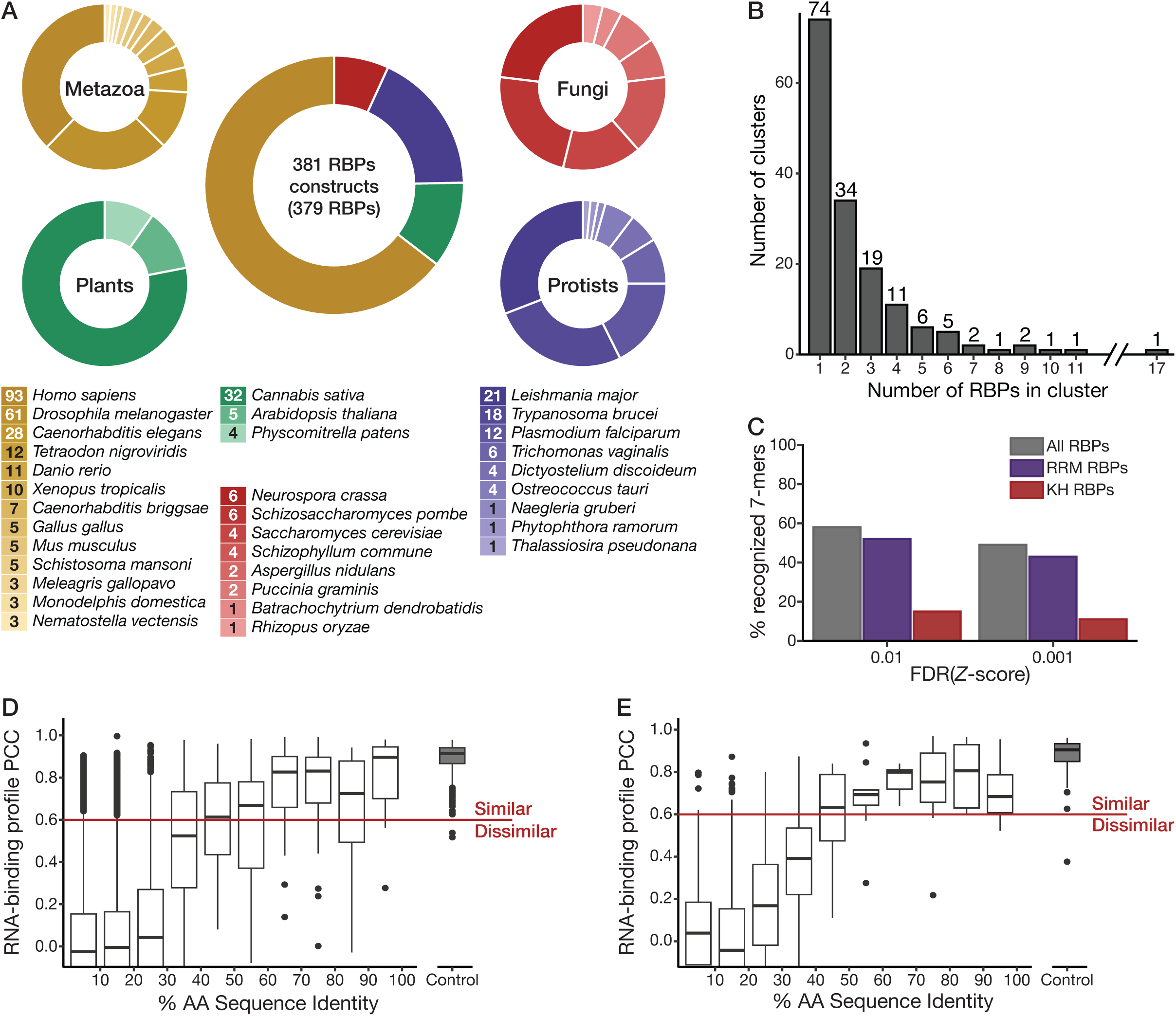
RNAcompete-measured RNA sequence specificities. **A**, The center donut chart depicts the breakdown of RNAcompete-measured RBPs across four eukaryotic kingdoms and 33 species, including both the RBPs measured for this study and those from Ray et al. 2013^1^. Clockwise from top left, donut charts depict the breakdown of RBPs by species for Metazoa, Fungi, Protists, and Plants. Legends adjacent to the donut charts show the number of measured RBPs for individual species. **B**, The RNAcompete-measured RBPs were split into clusters based on RNA-binding profile similarity; sequence specificities were hierarchically, agglomeratively clustered on the PCCs between RNA-binding profiles with complete linkage. Using a PCC cut-off of 0.6, 157 clusters were identified (**Table S1**) and the distribution of their sizes is displayed. **C**, Percentage of all 7-mers that are significantly bound (one-sided Z-test), FDR < 0.01 or < 0.001 (Benjamini-Hochberg FDR correction over the 16382 7-mers), by at least one RNAcompete-measured RBP, or at least one RNAcompete-measured RRM- or KH-domain RBP. **D**, **E**, Box plots show the distribution of RNA-binding profile PCCs for pairs of RBPs whose RBRs fall within the %AA ID range indicated on the x-axis. **D** depicts only the 308 RRM-domain-containing RBPs and **E** depicts only the 47 KH-domain-containing RBPs. As a control, the distribution of PCCs between RNAcompete Set A and Set B for the same experiment are displayed to the right.

**Figure S2.**
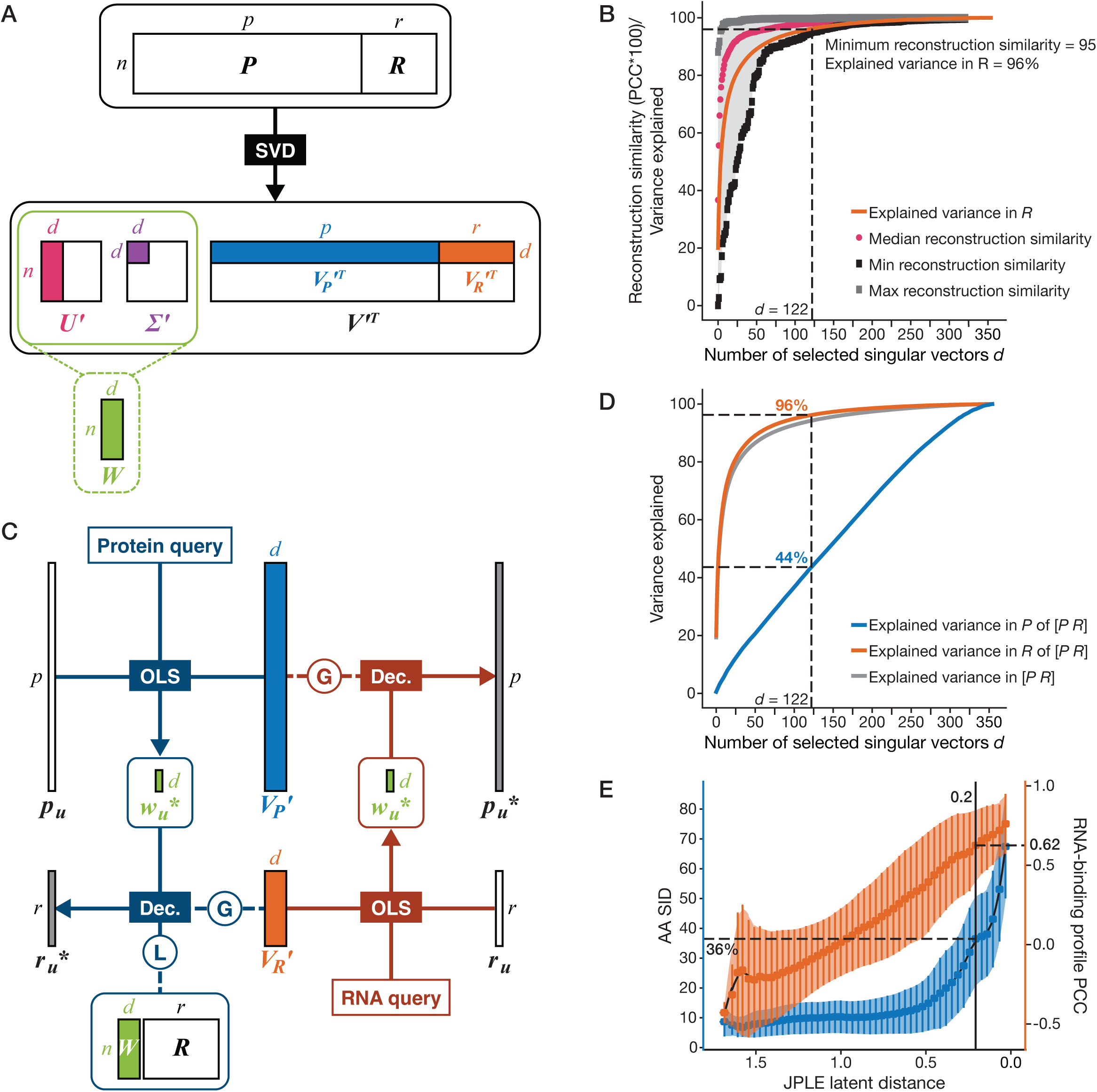
JPLE captures the association between amino acid sequence and RNA sequence specificity. **A,** Illustration of the JPLE training process for *n* RBPs. Singular value decomposition (SVD) is used to decompose the joint protein representation [***P R***] into ***U***, ***Σ***, and ***V^T^***. The *d* singular vectors and values contributing the most to the variance of ***R*** in [***P R***] are selected, leading to the submatrices ***U’***, ***Σ’***, and ***V’^T^***. The product ***W*** of ***U’*** and ***Σ’*** provides the *d*-dimensional latent embedding of the *n* RBPs. **B,** Distribution of the Pearson correlation coefficients (PCCs) between the reconstructed (***r\****) and measured (***r***) RNA-binding profiles (i.e., the reconstruction similarity), as a function of the number of maintained singular vectors *d*. The orange line represents the total fraction of variance explained in ***R*** of [***P R***]. The median, minimal, and maximal reconstruction similarities are displayed and the distribution is indicated in gray. To enable a minimum reconstruction PCC of 0.95 for all measured RBPs, *d* = 122 is required. **C,** Illustration of the JPLE inference process for RBPu. The left (in blue) showcases protein query, where the RBP’s latent embedding ***wu\**** is obtained by deconvolving its peptide profile ***pu*** into a mixture of the singular vectors in ***VP’***. Its RNA-binding profile ***r*u*\**** could be reconstructed through either global (labeled G) or local (labeled L) decoding. The right (in brown) showcases RNA query, where the RBP’s latent embedding ***wu\**** is obtained by deconvolving its RNA-binding profile ***ru*** into a mixture of the singular vectors in ***VR’***. Its peptide profile ***p*u*\**** could be reconstructed through global decoding. **D,** Variance explained in ***P*** and ***R*** of [***P R***], as a function of the number of selected singular vectors *d*. At *d* = 122, 44% and 96% of the total variance of ***P*** and ***R*** of [***P R***] are explained respectively. **E,** Relationship between RNA-binding profile PCCs and their JPLE latent distance (e-dist, i.e., cosine distance). JPLE was trained leaving clusters of RBPs with the same specificity (PCC>0.6) out, then embedding them into JPLE and measuring the distances between each other and to the RBPs in the training set. The latent distances between all RBP pairs were compared to their binding similarities (right y-axis) and the amino acid sequence identity (left y-axis). Lines and shaded area show smoothed mean and standard deviation across 50 equally sized bins. RBP pairs with < 0.20 latent distance possess an average RNA-binding similarity of 0.62 and AA SID of 37%.

**Figure S3.**
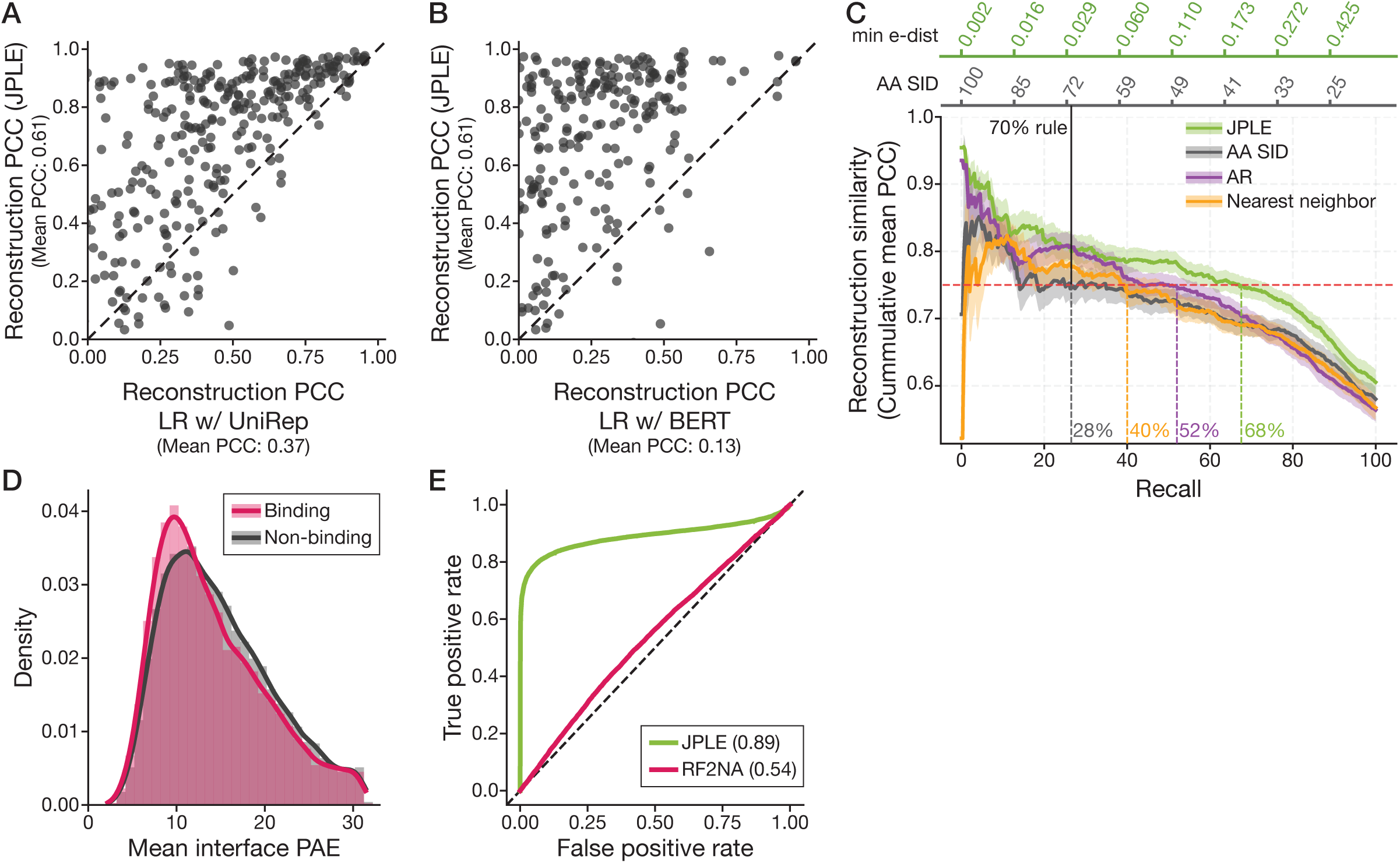
JPLE outperforms alternative methods. **A**, **B**, Comparison of the RNA-binding profile reconstructions generated by JPLE trained with 5-mer peptide features, to those generated by two Linear Regression (LR) models trained with deep learning features from TAPE: UniRep (**A**) and BERT (**B**). As in **Figure S2**, reconstruction PCCs are computed between the reconstructed (***r\****) and measured (***r***) RNA-binding profiles. **C,** Precision-recall curves for RNA-binding profile reconstructions generated by JPLE, amino acid sequence identity (AA SID), Affinity Regression (AR), and the nearest neighbor model (see Methods). Precision (y-axis) is the mean PCC for reconstructions at least as confident as the threshold (top axes). JPLE confidence is e-dist to the nearest neighbor; AA SID confidence is % amino acid identity; AR confidence is one minus PCC between the test and nearest neighbor’s embedding; the nearest neighbor model confidence is e-dist to the nearest neighbor. At the AA SID threshold of 70%, a mean PCC of 0.75 is achieved (red line). The recall for all four methods at a mean PCC of 0.75 is indicated. Standard error is shown in the shaded area around each line. **D**, Distribution of the mean interface predicted aligned errors (PAEs) for RoseTTAFold2NA’s predicted structures with high-affinity 7-mers (binding) and low-affinity 7-mers (non-binding) for all 355 RBPs. **E**, ROC curves compare the performance of RoseTTAFold2NA and JPLE in the task of differentiating high-affinity from low-affinity 7-mers. The predictions for RoseTTAFold2NA and JPLE are the mean interface PAEs (see **D**) and the predicted Z-scores on held-out RBPs, respectively. Numbers in brackets indicate the AUROCs.

**Figure S4.**
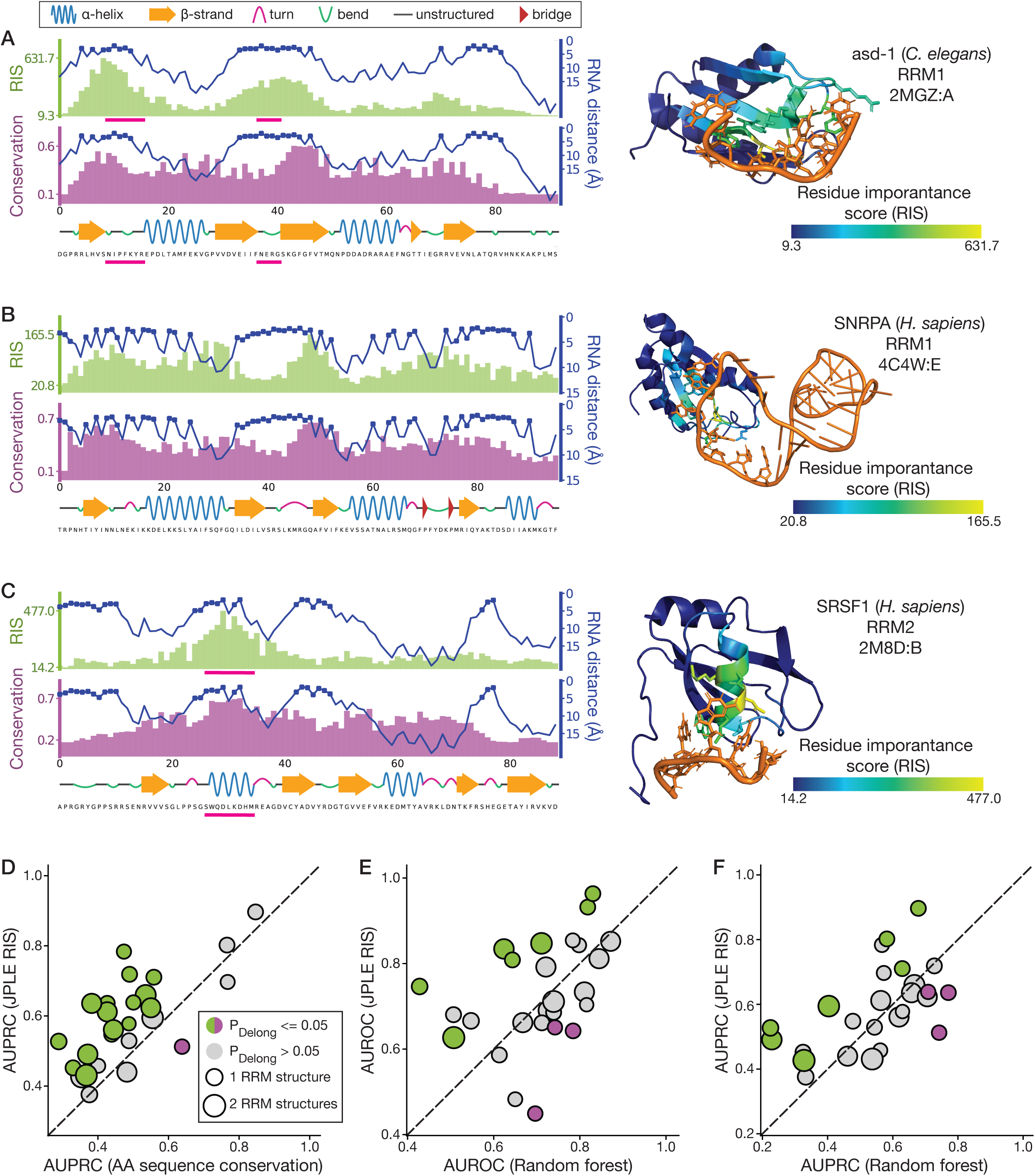
JPLE predicts RNA-interacting amino acids. **A-C**, The distance between individual residues and RNA (in Angstroms) is compared to JPLE residue importance scores (RISs) (top left) and conservation scores (middle left) for the RRM-RNA co-complex structure depicted to the right. A linear visualization of the protein secondary structure is depicted at the bottom left along with the protein sequence. RRM-RNA co-complex structures are coloured by JPLE RIS. The loops between β1 and α1 and between β2 and β3 found to confer specificity in RBFOX1^2^, the human homolog of *C. elegans* ASD-1, are indicated in **A** with a pink bar below the amino acid sequence and below the RIS histogram. Similarly indicated in **C** is the α-helix that confers sequence specificity in the depicted SRSF1 RRM^3^. **D**, Comparison between sequence conservation and JPLE RISs for predicting RRM domain interface residues, evaluated with AUPRC. Coloured circles indicate a significant difference in performance between the two scoring methods, as determined by the Delong test using AUROC values as in Figure 3C. **E-F**, Comparison between JPLE RIS and a random forest model trained using the following features: amino acid position specific sequence matrices from multiple sequence alignment, physico-chemical residue features, residue identity, conservation within a window of five amino acids. Results of evaluation by AUROC are shown in **E**, and by AUPRC in **F**. Points in both panels are coloured according to significance of the Delong test performed on AUROC values.

**Figure S5.**
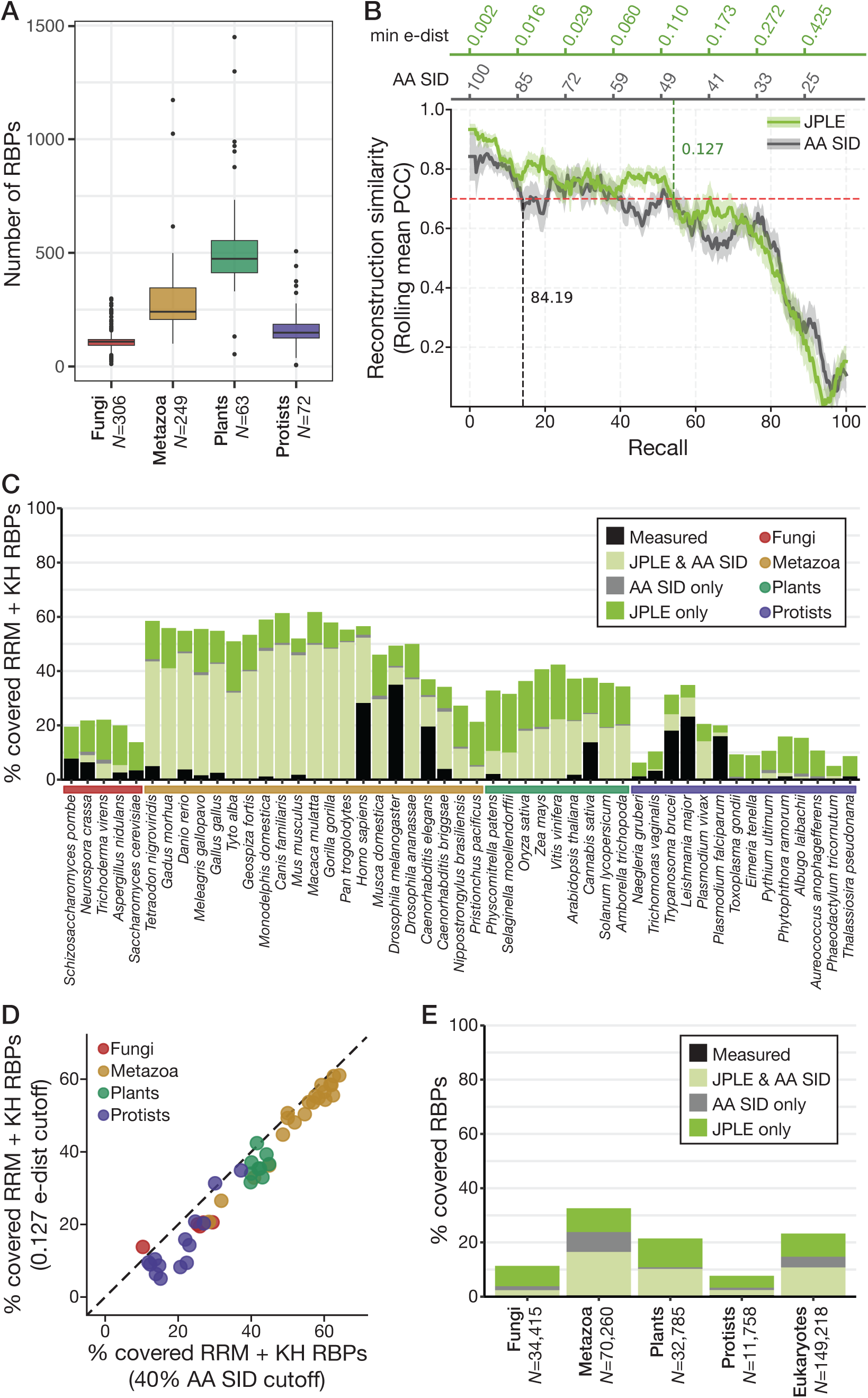
JPLE reconstructs RNA-binding specificities for thousands of eukaryotic RBPs. **A,** The number of RBPs identified in each of 690 eukaryotes split across four kingdoms. **B,** Precision-recall curves for RNA-binding profile reconstructions generated by amino acid sequence identity (AA SID) and JPLE. Precision (y-axis) is the mean rolling Pearson Correlation Coefficient (PCC) for reconstructions at least as confident as the threshold (top axes). The selection size for the rolling average window is 25 reconstructions. AA SID confidence is % amino acid identity, JPLE confidence is the minimum e-dist. Grey and green dashed lines indicate the confidence at which the rolling mean PCC first hits 0.70 (red dashed line). Standard error is shown in the shaded area around each line. **C,** The fraction of measured and reconstructed specificities for RRM- and KH-domain-containing RBPs for 49 representative species. The proportion of reconstructed specificities that were identified by AA SID, JPLE, or both are indicated. The kingdom to which the species belongs is indicated below the x-axis. **D**, Scatterplot displays the percentage of specificities for RRM- and KH-domain-containing RBPs that were reconstructed by JPLE (with an e-dist cutoff of 0.127) compared to a 40% AA SID for 49 representative species (listed in panel **C**). **E**, The fraction of measured and reconstructed RBP specificities for four eukaryotic kingdoms and all eukaryotes contained in EuPRI and on CisBP-RNA. This plot includes measured and reconstructed RBPs that do not contain RRM- or KH-domain-containing RBPs.

**Figure S6.**
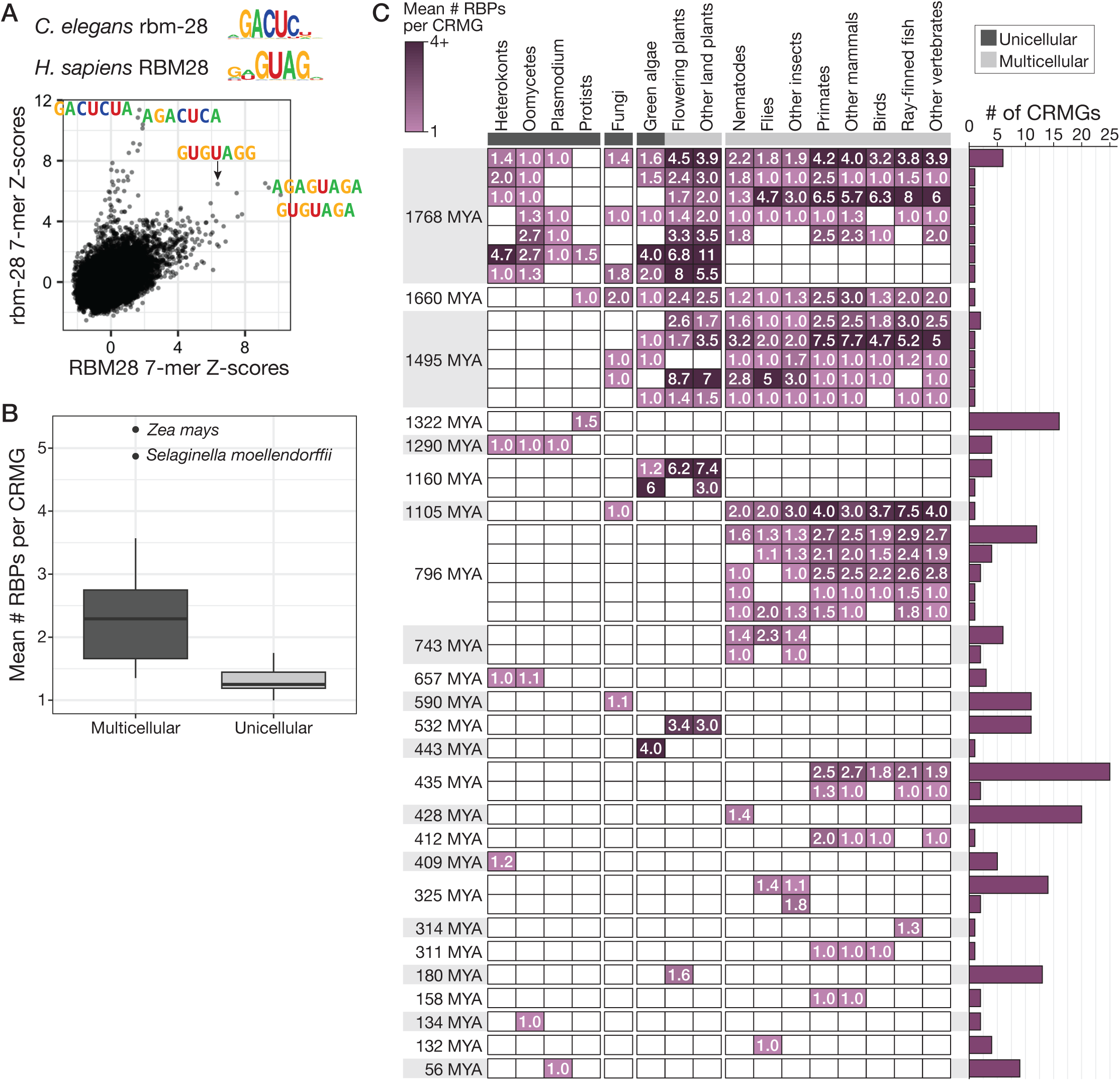
Evolution of eukaryotic CRMGs. **A**, RNAcompete 7-mer Z-scores are compared between human RBM28 and *C. elegans* ortholog *rbm-28*. RNAcompete motifs are shown above the plot, and some top 7-mers are directly labelled. **B**, The mean number of RBPs contained within a CRMG for multicellular (N=36) and unicellular (N=17) species. Only multi-species CRMGs containing an RBP with an e-dist of <0.2 to an RNAcompete-measured RBP were used for the calculations. Outlier species are labelled. **C**, The number of gained CRMGs at different time points are broken down by the major eukaryotic clades to which they belong. Cells in the heatmap are darkened to indicate the presence, in a given clade, of the CRMGs in the barplot to the right. The mean number of RBPs within the associated set of CRMGs in extant species belonging to the associated clade is displayed in each cell. The barplot on the right displays the number of CRMGs shared between the indicated clades that arose at the given time point. As in **B**, only multi-species CRMGs containing an RBP with an e-dist of <0.2 to an RNAcompete-measured RBP are shown.

**Figure S7.**
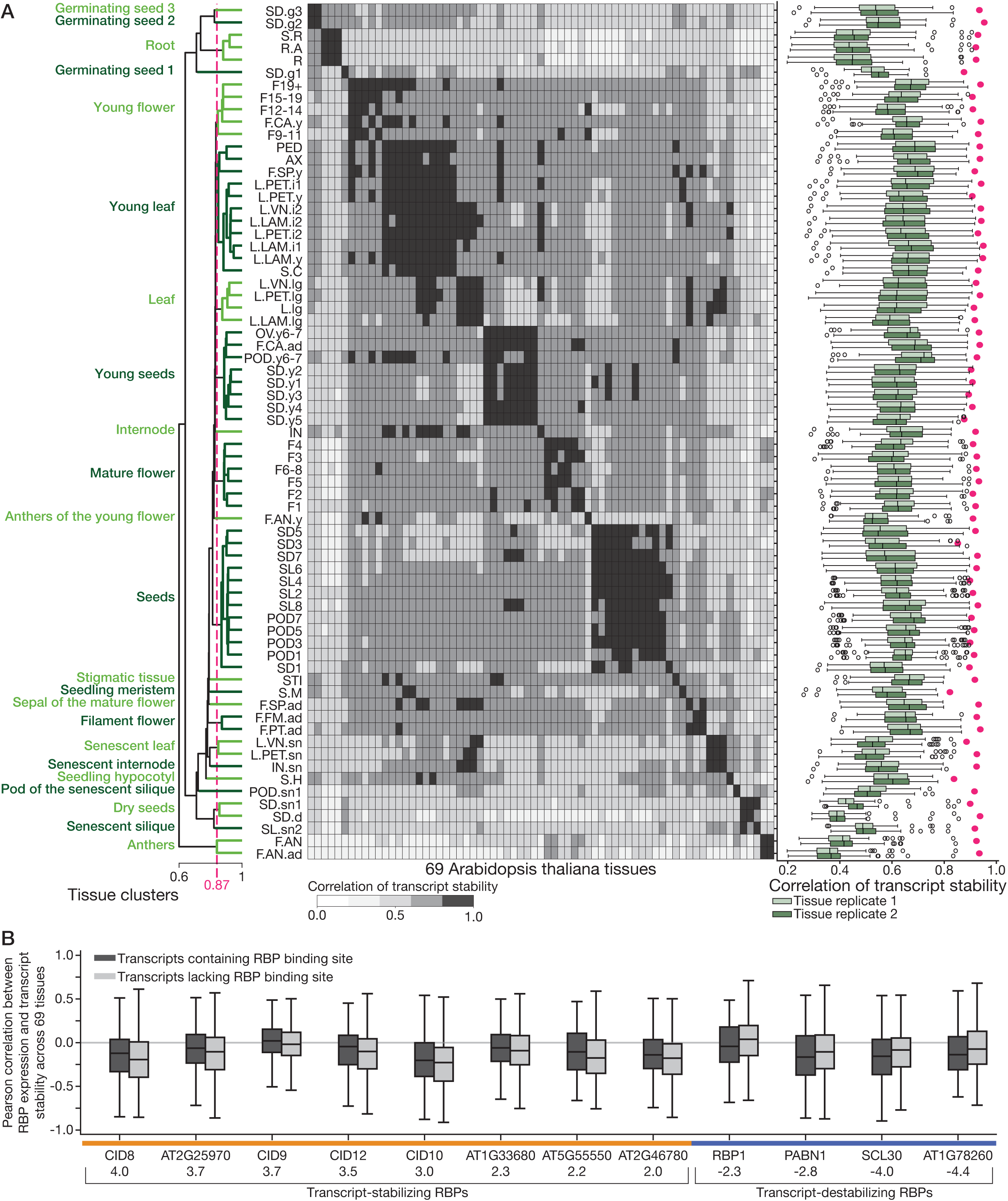
Predicting RNA stability regulators with reconstructed RNA sequence specificities in Arabidopsis thaliana. **A,** Heatmap displays the results of agglomerative clustering with single linkage on the Pearson Correlation Coefficients (PCCs) between the mRNA stability scores of the 6794 genes with reproducible scores. Tissue clusters were identified at a PCC cutoff of 0.87, generating 23 clusters, the names of which are indicated to the left and correspond to Figure 6B. To the right of the heatmap, boxplots display the mRNA stability score PCCs between each tissue replicate and all other tissue replicates, excluding the replicate from the same tissue. Tissue replicate correlations are displayed by the pink circle between each pair of boxes. **B,** Boxplot shows the distribution of PCCs between mRNA stability scores and RBP expression for the 12 RBPs identified as having a putative role in regulating stability.

